# Cell-cell communication as underlying principle governing color pattern formation in fishes

**DOI:** 10.1101/2025.08.21.671633

**Authors:** Marleen Klann, Saori Miura, Shu-Hua Lee, Stefano Davide Vianello, Robert Ross, Masakatsu Watanabe, Emma Gairin, Yipeng Liang, Harrison W. Hutto, Braedan M. McCluskey, Marcela Herrera, Lila Solnica-Krezel, Laurence Besseau, Simone Pigolotti, David M. Parichy, Masato Kinoshita, Vincent Laudet

## Abstract

The diverse pigmentation patterns of animals are crucial for predation avoidance and behavioral display, yet mechanisms underlying this diversity remain poorly understood. In zebrafish, Turing models have been proposed to explain stripe patterns, but it is unclear if they apply to other fishes. In anemonefish (*Amphiprion ocellaris)*, we identified *gja5b*, a gene orthologous to zebrafish *leopard* and encoding a connexin involved in pigment cell communication, as responsible for the *Snowflake* phenotype. Using CRISPR/Cas9 and transgenesis, we recapitulate the *Snowflake* phenotype and show expression of *gja5b* in iridophores. A matching allele was recovered in zebrafish, revealing complementary requirements in both species. Our findings highlight conserved roles of gap junction mediated communication in pigment patterning across divergent teleosts.

## Introduction

Pigment patterns are among the most striking traits in animals, playing a role in predator avoidance, communication, and thermoregulation (*1*). Understanding how pigment patterns form and diversify is therefore crucial in elucidating the processes underlying morphological organization. In ectothermic vertebrates, these patterns emerge from the spatial arrangement of chromatophores – specialized pigment cells (such as black melanophores and yellow/orange xanthophores) or light reflecting cells (like iridescent/white iridophores) in the skin.

Studies of zebrafish (*Danio rerio*, Cyprinidae) reveal how self-organizing interactions of chromatophores are coordinated by morphogenetic behaviors, specification and differentiation to generate and maintain an organized pattern (reviewed in (*2*)). Theoretical and empirical investigations in this model species have shown that the dynamics of stripe formation can be described with Turing-type reaction-diffusion models, in which short-range activation and long-range inhibition between pigment cells, in particular via gap junctions, are central in establishing patterns (*3, 4*). This is illustrated by stripe formation failure in spotted *leopard* and *luchs* mutants. These harbor mutations in *gap junction gene 5b* (*gja5b*; *connexin 41*.*8*) and *gja4* (*connexin 39*.*4*), respectively, encoding Connexins that function in gap junctions of melanophores and xanthophores (*5, 6*). It is not known if similar principles explain the wide range of patterns across in other teleost fish lineages (*4, 7, 8*). For instance, the distantly related anemonefish *Amphiprion ocellaris* (Pomacentridae; Fig. 1A), has a simple and robust color pattern: an orange body adorned with three vertical black bordered white bars, which appear sequentially from head to tail during metamorphosis (*9*). This color pattern remains unchanged as the fish grows, raising the possibility that it develops by mechanism independent of a Turing process (e.g., positional cues (*9*)).

**Fig. 1.**
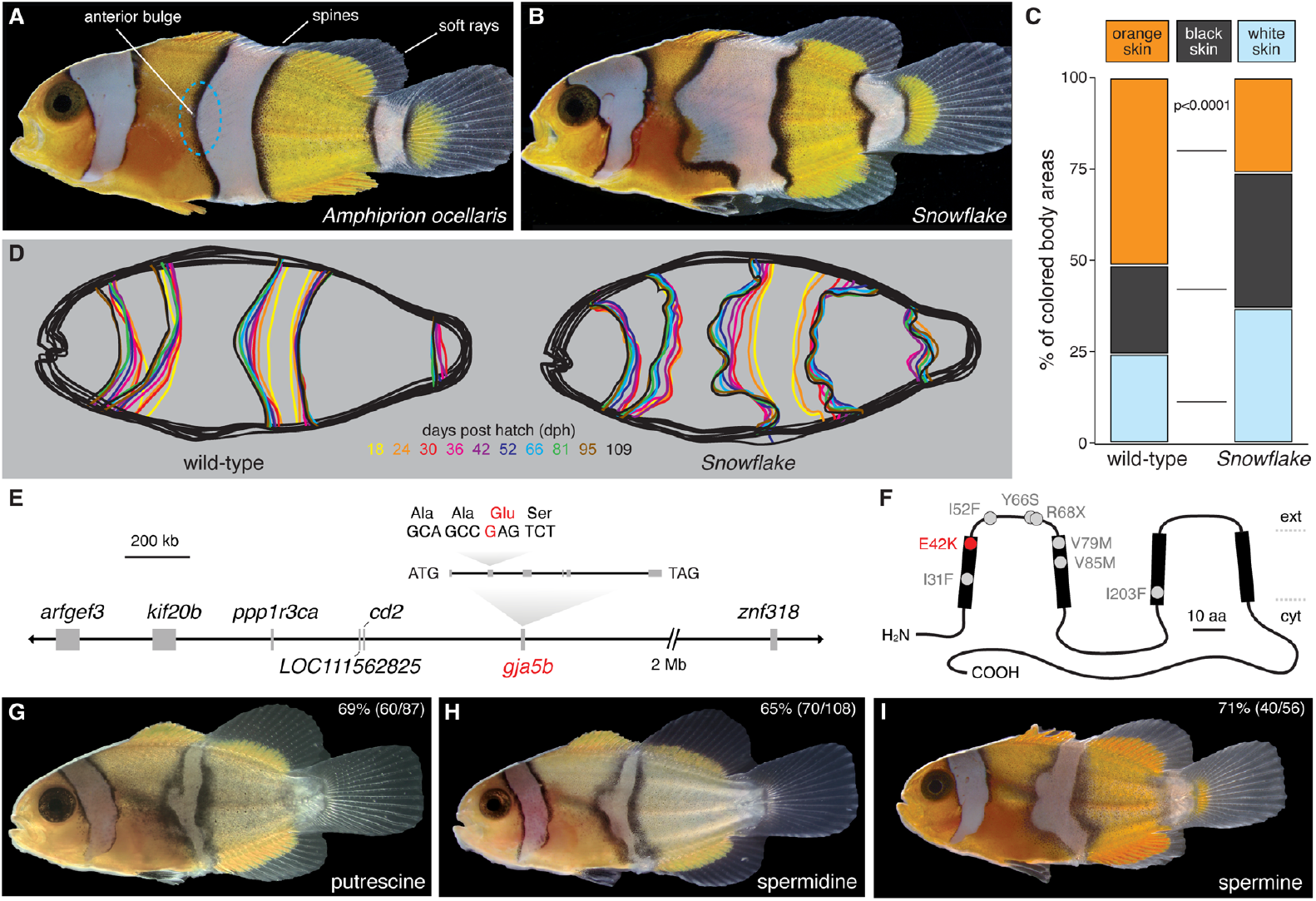
*Snowflake* phenotype, underlying gene candidates, and pharmacological gap junction inhibition. **(A**) The wild-type *Amphiprion ocellaris* color pattern comprises three vertical white bars with black edges on an orange body. The trunk bar shows an anterior bulge (blue oval). The dorsal fin spines and rays are indicated. (**B**) The *Snowflake* mutant color pattern is characterized by increased white areas with undulating, thicker outlines. (**C**) Comparison of colored body areas between wild-type and *Snowflake*, with increased white and black skin in *Snowflake* (overall MANOVA, F_3,54_=78, overall and all individual comparisons, p<0.0001; n=23 wild-type, 35 *Snowflake*). (**D**) During color pattern development the *Snowflake* phenotype deviates from the wild-type from the beginning with broader white areas that develop increasingly jagged boundaries as development proceeds (dph, days post-hatching). (**E**) GWAS analysis revealed 7 potential candidate genes, of which only one, *gja5b*, exhibited a coding sequence variant fully associated with the *Snowflake* phenotype. (**F**) Gja5b showing transmembrane domains (thick line segments), E42K mutation of *Snowflake* and previously identified variants of zebrafish. (ext, extracellular face; cyt, cytosolic face). (**G** to **I**) When pre-metamorphic wild-type larvae were treated with polyamine gap junction inhibitors, putrescine (G), spermidine (H), and spermine (I), fish developed irregular bars, similar to *Snowflake*. These abnormal pigment patterns arose regardless of charge (+2, +3, +4, respectively), suggesting the blocking mechanism is highly sensitive.

Here, making use of one of many spontaneously arisen variants of anemonefish bred for tropical fish hobbyists (*10*), we focused on the *Snowflake* mutant (Fig. 1B), which exhibits jagged boundaries between colored bars that suggest defects in chromatophore communication. We show that a point mutation in *gja5b* underlies the *Snowflake* phenotype, thereby demonstrating a role for this locus and gap junction interactions in anemonefish and zebrafish, which last shared a common ancestor more than 200 million years ago (*11*). Nevertheless, anemonefish *gja5b* is expressed by iridophores, unlike zebrafish *gja5b*, and the *Snowflake* phenotype is qualitatively different from that of an identical mutation in zebrafish. Lastly, by engineering a parallel cellular context in zebrafish and modulating gap junctional communication with iridophores, we find defects in pattern formation consistent with those of *Snowflake*. Our findings illustrate how the same molecular players used in different contexts contribute to phenotype diversification and underscore cell-cell communication as a key principle in pattern formation.

## Results

### *Snowflake* mutants are defective for pattern boundary positioning

In comparison to wild-type *A. ocellaris* (Fig. 1A), the color pattern of *Snowflake* (Fig. 1B) differs in three ways (see Supplementary text): (1) white areas are increased (Fig. 1C), (2) more melanophores occur within black edges (Fig. 1C), and (3) bar outlines are uneven (Fig. 1D). Interestingly, every *Snowflake* fish has a unique appearance – the ways by which the black edge is curved and uneven differs between individuals. Yet each fish shows a very high degree of bilateral symmetry (fig. S1A-C).

Anemonefish exhibit two distinct color patterns during development. Larval coloration is dominated by two horizontal stripes of black melanophores. At this stage, wild-type and *Snowflake* fish were phenotypically indistinguishable (fig. S1D). Upon metamorphosis, when the adult color pattern starts appearing, the *Snowflake* pattern became characterized by broader white areas (yellow lines in Fig. 1D). The mutant pattern was fully established after 2-3 months, with only minor alterations during later growth, indicating a defect in pattern formation rather than maintenance.

### A connexin gene, *gja5b*, is mutated in *Snowflake*

Using GWAS and subsequent single nucleotide variant detection (see Supplemental text), seven linked genes on chromosome 16 were identified as plausible candidates for *Snowflake* (Fig. 1E, fig. S2A). Among these genes, *gap junction protein alpha 5b (gja5b*), is the anemonefish orthologue (fig. S2B) of zebrafish *leopard*, encoding gap junction protein Connexin 41.8 (Cx41.8) (*5*), a major actor in the Turing-like patterning system in zebrafish (*12*). Using Sanger sequencing on *Snowflake* fish from different aquaculture origins, we ruled out all genes other than *gja5b* (table S1). In *Snowflake*, we identified a 124G>A transition mutation, leading to a Glu42Lys (E42K) amino acid substitution within the extracellular side of the first transmembrane domain of the Connexin (Fig. 1F). We observed that all *Snowflake* mutants surviving past the first three days of larval life were heterozygous, while homozygous *Snowflake* mutants developed and hatched but died rapidly after without any overt phenotype (fig. S1D). Consistent with a critical role of *gja5b* during normal early larval development, stage-wise transcriptomic data revealed an upsurge in its expression at 2 days post-hatching (stage 2 in fig. S2C).

To further test the correspondence of *Snowflake* and *gja5b*, we used two independent approaches. First, given that the candidate gene encoded a gap junction protein, we predicted that pharmacological inhibition of gap junctional communication in wild-type fish should lead to defects resembling those of *Snowflake*. We found that each of three polyamines (putrescine, spermidine, and spermine) known to inhibit gap junction communication (*13*) resulted in uneven bar outlines strikingly similar to those of *Snowflake* in genotypically wild-type fish (Fig. 1G–I). By contrast, treatment of *Snowflake* individuals with polyamine did not alter the mutant phenotype (fig. S2D), as anticipated if the mutation already blocks gap junctional communication. Second, we predicted that if the *Snowflake* phenotype is due entirely to the observed mutation, rather than regulatory or other variants, then a corresponding mutation generated independently in an otherwise wild-type background should phenocopy *Snowflake*. Accordingly, we used CRISPR/Cas9 homology directed repair (HDR) to replace exon 4 of *gja5b* in wild-type fish with a homologous sequence having an E42K mutation (fig. S2E,F). In G0 fish, 14% of 102 surviving juveniles exhibited abnormal color patterns reminiscent of *Snowflake* (Fig. 2A), with individual variation and asymmetries (distinct from the normal bilateral symmetry) as expected given genetic mosaicism (compare Fig. 2A and 2B, two sides of the same fish). We additionally targeted the E42 codon directly by CRISPR/Cas9, to induce a spectrum of mutations by non-homologous end joining. Survival rates at hatching (21%) and for juveniles (10%) were reduced in comparison to injections replacing the entire exon (49% at hatching and 26% for juveniles), consistent with a subset of biallelic mutations leading to early lethality, as observed in *Snowflake* homozygotes. Nevertheless, all 13 surviving juvenile fish displayed *Snowflake*-like phenotypes, with most fish showing an effect on all three white bars (Fig. 2C,D). Together these findings indicate that *Snowflake* results from an E42K substitution in *gja5b*, the gene encoding gap junction protein Connexin 41.8.

**Fig. 2.**
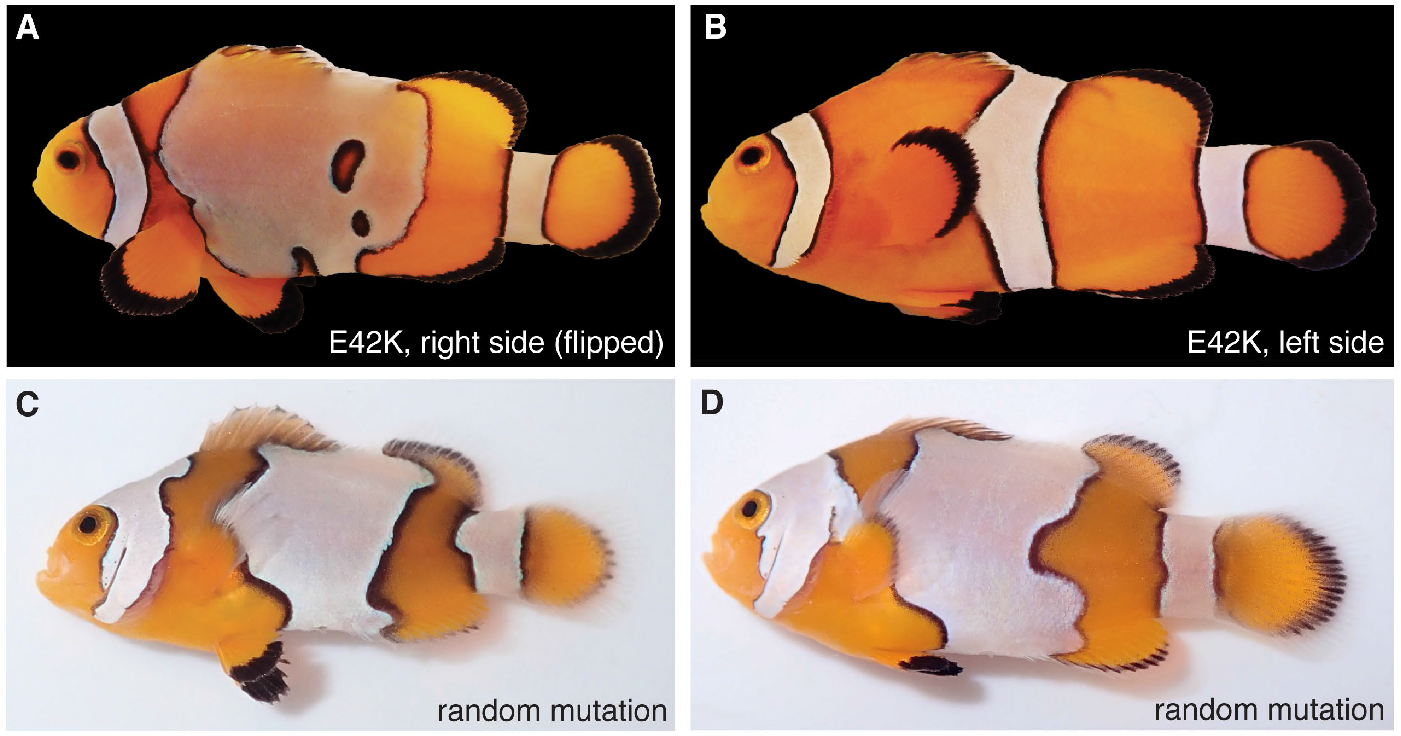
CRISPR/Cas9 genome editing replicates the *Snowflake* phenotype. (**A** and **B**) Targeting wild-type *gja5b* with an E42K substitution resulted in *Snowflake*-like phenotypes (A, severe bar disturbance; B, opposite side of the same mosaic individual with wild-type pattern). (**C** and **D**) CRISPR/Cas9 mutations induced by NHEJ in the vicinity of E42 also led to phenotypes similar to *Snowflake* in G0 fish.

### Cell-cell communication governs color pattern formation in anemonefish

To further understand the function of gap junctional communication during color patterning in anemonefish, we assessed the expression of *gja5b*. RNA sequencing from wild-type *A. ocellaris* (WT) and the *Black* mutant (BO) (*10*), used to obtain sufficient quantities of black scales, revealed high levels of *gja5b* transcript in white bars, enriched in iridophores (Fig. 3A). *gja5b* transcript was found only at low levels in orange scales (enriched in xanthophores) and black scales (enriched in melanophores). The finding that *gja5b* is predominantly expressed in iridophores was surprising as *gja5b* of zebrafish is expressed primarily in xanthophores and melanophores (fig. S3A). To confirm the spatial expression pattern of *gja5b*, we produced transgenic fish harboring GFP driven by *gja5b* promoter using the transposon system (fig. S3B). In 18 of 19 (95%) G0 mosaic fish, we detected extensive expression of EGFP in white bars along with other cell types (Fig. 3B; fig S3C). These findings suggest that, in anemonefish, *gja5b* primarily contributes to pigment patterning through its role in iridophores (see Supplemental text).

**Fig. 3.**
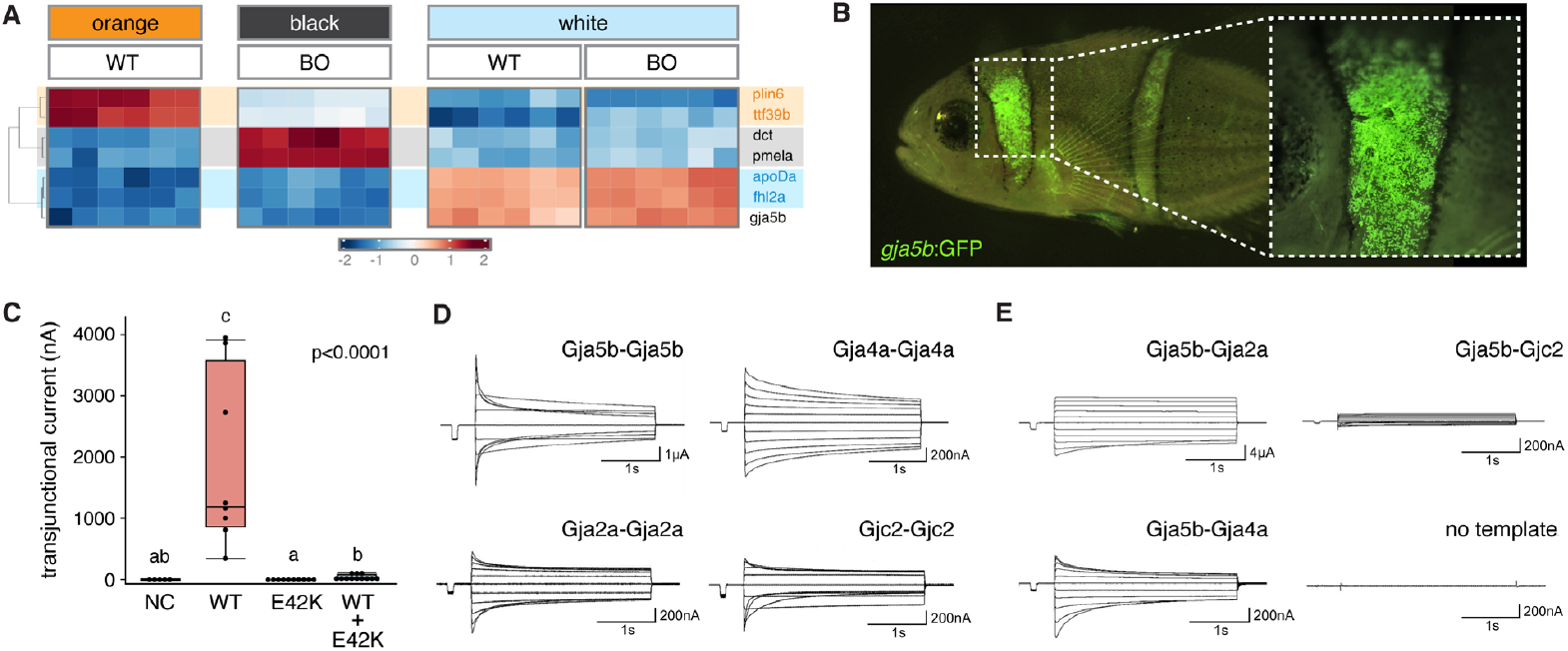
Anemonefish *gja5b* expression and transjunctional current assays. (**A**) RNA-sequencing of scale samples enriched for orange xanthophores (marker genes: *plin 6* and *tt39b*), black melanophores (marker genes: *dct* and *pmela*) and white iridophores (marker genes: *apoDa* and *fhl2a*) revealed highest *gja5b* expression in association with iridophores. Scales were collected from wild-type fish (WT) and *Black A. ocellaris* mutant (BO). (**B**) The expression of *gja5b* was monitored by GFP. The GFP gene driven by *gja5b* promoter was introduced into wild-type fish (for construct see fig. S3B). Predominant GFP expression was found in white bar areas. (**C**,**D**,**E**) Transjunctional current assays in *Xenopus* oocytes were performed to test the formation and functionality of gap junctions. (C) In comparison to transmitted current at -100mV for wild-type (WT) Gja5b, little to no current was transmitted by E42K-mutated *gja5b*, singly or in combination with wild-type. NC, no-construct control. (Overall Kruskal-Wallis, χ^2^ approximation=24.6, p<0.0001; shared letters over groups indicate medians not significantly different from one another by *post hoc* Steel-Dwass comparisons.) (D). Homotypic gap junction of Gja5b, Gja2a, Gja4a, and Gjc2 are functionally active. (E) Heterotypic gap junctions between Gja5b and Gja2a and between Gja5b and Gja4a are functionally active, while gap junctions between Gja5b and Gjc2 appear to either fail or have extremely low activity.

The heterozygous phenotype of E42K substitution raised the possibility of dominant negative activity, as inferred previously for some semi-dominant alleles of zebrafish (*14*). We therefore assayed the function of E42K alleles in a *Xenopus* oocyte transjunctional current assay (*15*). Injected wild-type *gja5b* cRNA allowed transjunctional currents, demonstrating that gap junctions were formed and electrical signals propagated (Fig. 3C). By contrast, E42K cRNA abolished current, indicating a failure in electrical signal propagation (Fig. 3C). Co-injection of equimolar E42K and wild-type cRNAs further indicated a strong dominant negative effect (Fig. 3C).

In zebrafish Gja5b forms heteromeric gap junctions with Gja4 (mutated in *luchs* (*16*)) and we therefore searched candidate partner connexins for anemonefish Gja5b. In *A. ocellaris* 42 gap junction genes could be identified (fig. S4A). We screened expression of these gap junction genes in scales of the three different colors of wild-type fish, detecting 23 (fig. S4A, highlighted). Most gap junction genes (18 out of 23) were expressed in two or more color-regions whereas *gja5a* and *gja5b* were expressed predominantly in the white area (fig. S4B). Interestingly, *gja4a* was expressed in orange xanthophore-enriched tissue and black melanophore-enriched tissue consistent with the expression of its orthologue in zebrafish. In anemonefish, iridophores and xanthophores are not in direct contact (*17*) and therefore gap junction formation is unlikely, but it is possible that Gja5b of iridophores forms gap junctions with Gja4a of melanophores. We therefore tested if three connexins consistently expressed across samples by xanthophores and melanophores (Gja2a, Gja4a, and Gjc2; fig. S4B, asterisks, figS4C) can form heterotypic gap junctions with Gja5b expressed predominantly by iridophores. Injection of cRNAs for each of the three connexins into *Xenopus* oocytes showed that all of them could form functional gap junctions on their own (Fig. 3D). We then performed assays using pairs of oocytes, one injected with Gja5b and the other with one of the three connexins. When Gja2a or Gja4a was injected, functional gap junctions were formed with Gja5b-injected oocytes (n=3 for Gja2a, n=6 for Gja4a; Fig. 3E). In contrast, heterotypic gap junctions formed between Gja5b and Gjc2 either failed to produce functional gap junctions or showed extremely low activity consistent with the absence of the ExxxE domain at position 12-16 in its protein sequence (n=4; Fig. 3E).

### Iridophore-specific manipulation of *gja5b* disrupts pigment patterning in zebrafish

Expression of *gja5b* in white bars of *A. ocellaris* and its presumed function in iridophores contrasted with zebrafish, in which *gja5b* is expressed and required in melanophores and xanthophores. Zebrafish *gja5b* mutant alleles confer a range of spotted phenotypes with null alleles being recessive and homozygous viable (*5, 16, 18*). Fortuitously, we identified a new ethylnitrosourea-induced allele in zebrafish, *gja5b*^*stl710*^, which presents the identical nucleotide change and amino acid substitution (E42K) as *Snowflake*, allowing a direct comparison of phenotypes between the two species. Zebrafish *gja5b*^*stl710*^ exhibited a heterozygous phenotype of irregular stripes grading into spots and a homozygous viable phenotype of few widely dispersed melanophores associated with a failure to coalesce into stripes and severe attenuation of electrical current flow (Fig. 4A; fig. S5A,B; movies S1,2).

**Fig. 4.**
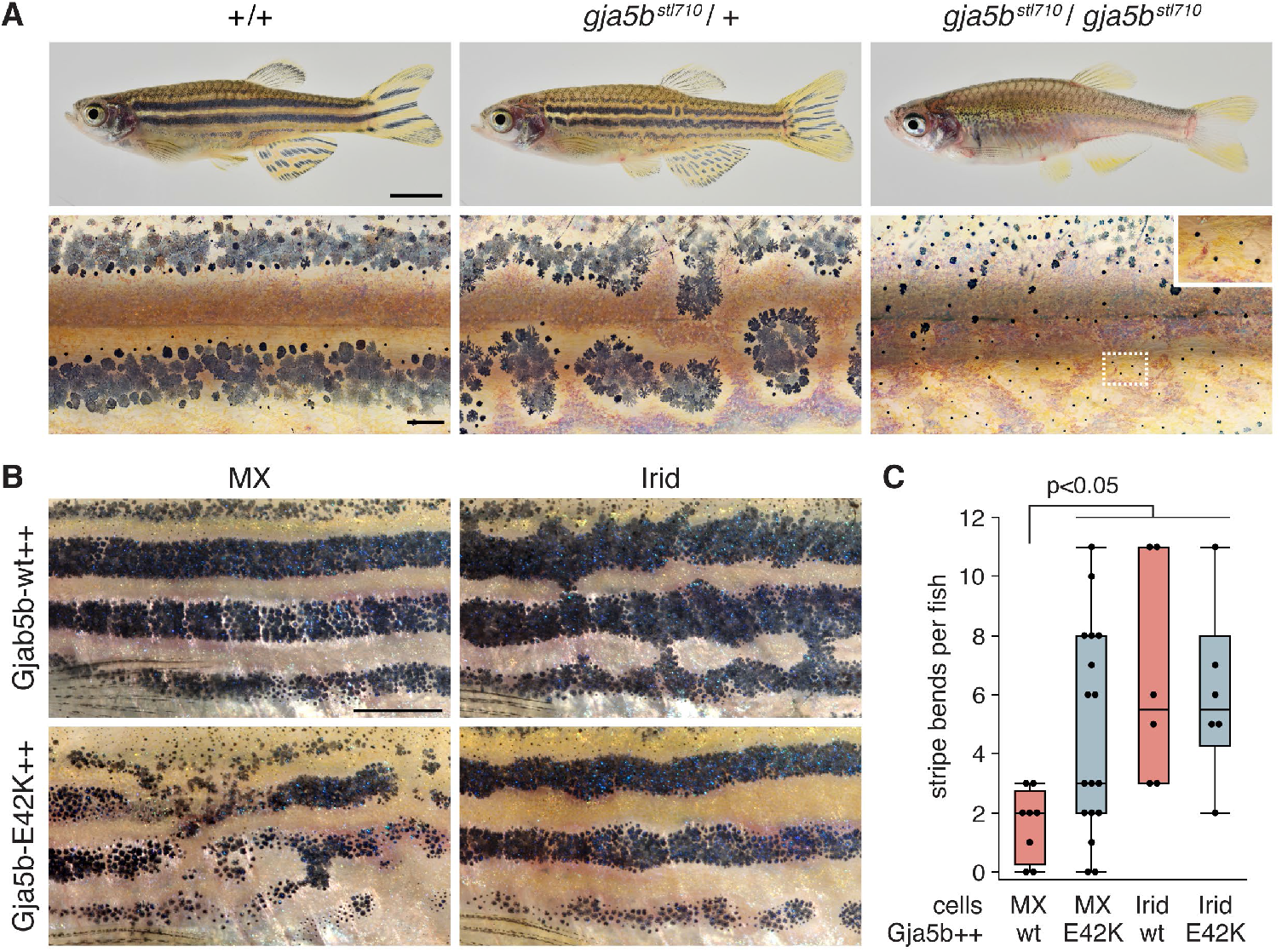
An E42K mutation in zebrafish and patterning consequences of gap junctional perturbation by misexpression. (**A**) Comparison of wild-type zebrafish and mutants heterozygous or homozygous for E42K mutant allele *gja5b*^*stl710*^. Inset shows higher magnification of melanin in individual, dispersed melanophores of homozygous mutant. (**B**) Overexpression of wild-type *gja5b* (*gja5b*-wt++) in melanophores and xanthophores (MX) did not result in a visible phenotype (top left) whereas overexpression of mutant Gja5b (Gja5b-E42K++) led to phenotypes similar to *leopard* mosaics (bottom left). By contrast wild-type Gja5b expressed by iridophores (Irid) resulted in melanophores trespassing into interstripes (top right), and expression of mutant Gja5b resulted in wandering stripe boundaries and expanded interstripe regions (bottom right). (**C**) Pattern irregularities assessed as bends in stripe boundaries were significantly increased when iridophores expressed wild-type Gja5b ectopically, or when either set of chromatophores expressed E42K. Scale bars, 1 cm (in A, upper), 200 μm (in A, lower), 1 cm (in B).

Given the correspondence between these mutations and their consequences for gap junctional communication, and considering the expression and presumed function of *gja5b* in iridophores of anemonefish, we hypothesized that gap junctions of zebrafish iridophores – in addition to those of melanophores and xanthophores – function in stripe formation. Indeed, iridophores express several gap junction genes, including low levels of *gja5b* during adult stripe development (fig. S3A). If iridophores participate in gap junctional communication, we predicted that blockade of such communication should lead to pattern defects. We therefore mosaically expressed wild-type or dominant negative Gja5b^E42K^ in wild-type zebrafish using the promoter of *defbl1* which drives expression at high levels in iridophores but not melanophores or xanthophores (*19*). Overexpression of Gja5b^WT^ in iridophores led to melanophores trespassing into light interstripes, whereas Gja5b^E42K^ led to expanded interstripes with irregular boundaries (Fig. 4B,C; fig. S5C). These phenotypes were specific to iridophores, as overexpression of Gja5b^WT^ specifically in melanophores and xanthophores using a promoter derived from *mitfa* (*20*) did not affect stripe borders whereas expression of Gja5b^E42K^ in these cells led to phenotypes resembling those of individuals chimeric for *leopard* mutations (*18*) (Fig. 4B).

These results support a model in which all three major chromatophore classes are responsive to changes in gap junctional activities, with these activities contributing to the formation of the very diverse color patterns of teleost fish species as phylogenetically distant as zebrafish and anemonefish, emphasizing the importance of gap junction mediated communication in pigment patterning.

### Boundary modeling reveals low tension and dynamic noise in shaping *Snowflake* phenotype

We turned to biophysical modeling to better understand the effect of the *Snowflake* mutation on white bar patterning. We implemented an approach based on the Edwards-Wilkinson model, which describes the dynamics of membranes and interfaces (*21*). This approach assumes that the boundary separating color patches develops under the effect of random fluctuations and surface tension. At variance with traditional biochemical models (*22*), this approach does not require detailed knowledge of gene regulatory networks and cell-cell interactions. It is therefore particularly appropriate for anemonefish, an emerging model organism for which our knowledge of the processes is still limited (*7*).

To apply the model to anemonefish patterning, we traced boundary shapes of the trunk bar (both anterior and posterior individually) in developmental series of wild-type and *Snowflake* (Fig. 5A). We then applied a band-pass filter to the data to remove large scale fluctuations due to body shape and therefore not captured in the model, as well as small-scale fluctuations due to experimental error in measuring contours (Fig 5B and Methods). Resulting estimates of boundary fluctuations in wild-type and *Snowflake* were in excellent agreement with the predictions of the Edwards-Wilkinson model (fig. S6). In this comparison, the K ratio, representing the ratio between the intensity of random fluctuations and surface tension, was about one order of magnitude larger in *Snowflake* than in wild-type (K = 0.0039±0.0001 for WT anterior; K = 0.00426±0.00005 for WT posterior; K = 0.082±0.005 for SF anterior; K = 0.13±0.01 for SF posterior).

**Fig. 5.**
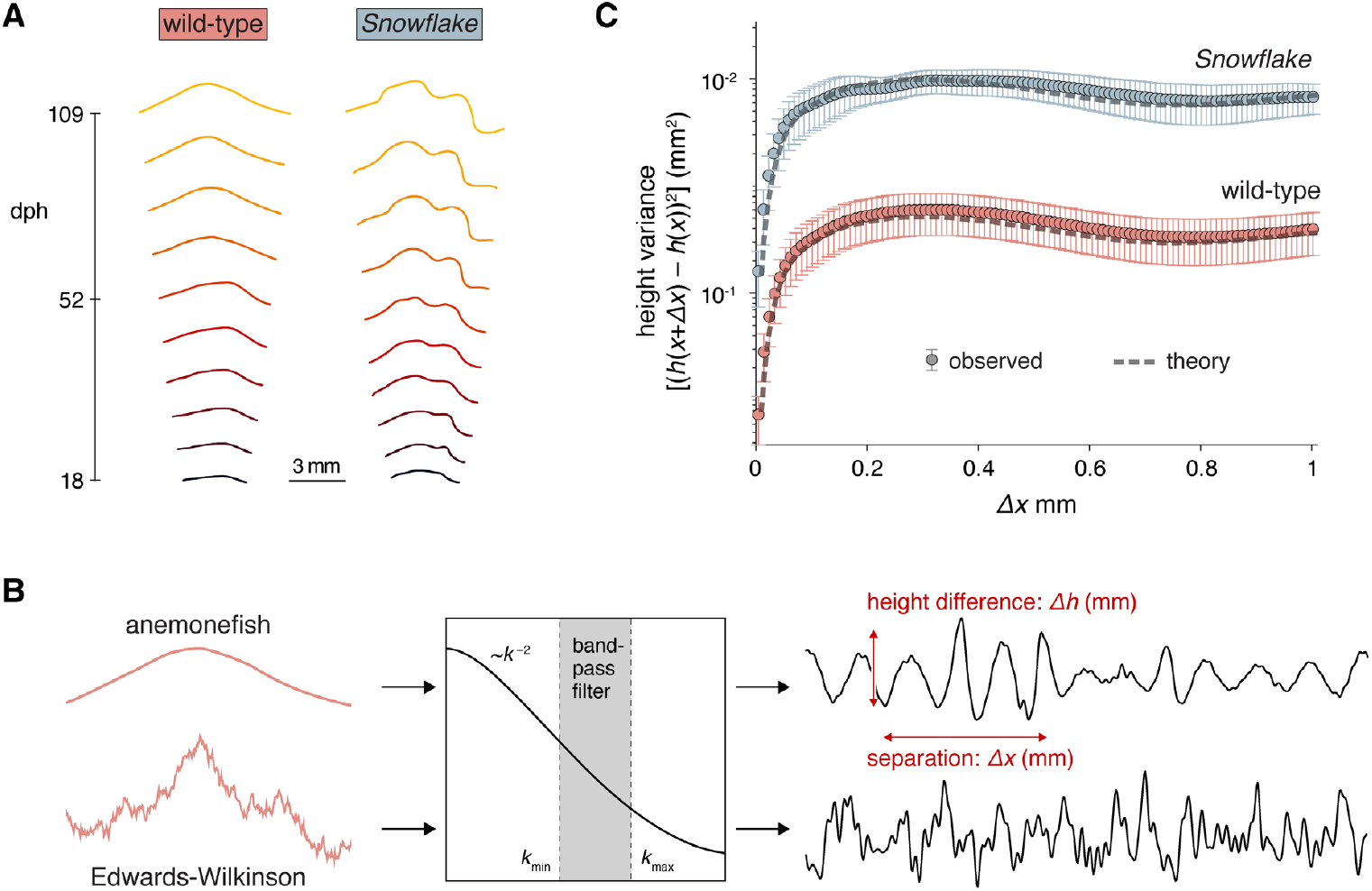
Boundary modeling in wild-type and *Snowflake* fish. (**A**) Examples of boundary shapes traced in developmental series of wild-type (WT) and *Snowflake* (SF). (**B**) Representation of selected modeling steps applied during Edwards-Wilkinson. (**C**) Theoretical prediction of the Edwards-Wilkinson model and actual measurements fitted very well.

We used the model to predict boundary roughness quantified as the variance of the boundary height difference *Δ*h between points on the line at a certain distance *Δ*x (Fig. 5B and Methods). According to the model, the roughness increased approximately linearly with distances on the order of 0.2 - 0.3 mm, and then plateaued at an approximately constant value, which is dependent on the parameter K. This theoretical prediction agreed with the boundary roughness measured directly from experimental data, both for the anterior trunk bar boundary (Fig. 5C) as well as the posterior trunk bar boundary (fig. S6E). The comparison of observation and modelling supports that both the anterior and posterior boundaries of the trunk bar in *Snowflake* are rougher, as a combined effect of more intense fluctuations and reduced surface tension.

## Discussion

In this study, we identified *gja5b* as the gene mutated in *Snowflake*, showing that a single amino acid change in the encoded connexin Cx41.8 disrupts pigment pattern formation in anemonefish. The *Snowflake* phenotype, with its uneven and curved bar outlines, emerges from a defect in boundary positioning during metamorphosis. Modeling revealed that these jagged contours likely reflect higher dynamic noise and/or reduced tension between color domains. The fact that each *Snowflake* fish looks different, but the two sides of each fish appear strikingly similar leads us to believe that contributions of random noise are limited and much of the phenotype results from reduced tension. Biologically, rather than affecting the differentiation of specific pigment cell classes, the mutation disturbs how these cells communicate and determine their respective positioning. In wild-type fish, sharp boundaries likely rely on precise communication between iridophores and melanophores, a process compromised in *Snowflake* due to the *gja5b* mutation, resulting in rougher outlines and broader white areas. Interestingly, expressing the same mutant allele in zebrafish iridophores also caused boundary defects, highlighting a conserved role for gap junction dependent interactions despite differences in cell-type specific expression of *gja5b* itself. Our data suggests that cell-cell communication is not only critical for boundary shaping, but also for maintaining the proper chromatophore ratios during pattern formation as shown by the broader white areas in *Snowflake*.

The roles of gap junctions in pigment patterning appear to be both widespread and highly adaptable across teleost fish. In our study, the E42K mutation in the connexin *gja5b* was shown to disturb pigment cell arrangement in both anemonefish and zebrafish, even though the gene is expressed in different chromatophore types between the two species. Studies on species as diverse as marble trout, chicken, and quail also highlighted the importance of *gja5* orthologues in color patterning (*23-25*). These findings indicate that connexins, especially *gja5* but likely others too, have been co-opted repeatedly for color pattern formation across distant teleost fish and in other vertebrates. What differs is not necessarily the function of the protein itself, but the context in which it was deployed: the pigment cell types, their spatial arrangement, and their interaction dynamics. This flexibility suggests that pigment pattern evolution can proceed not only through changes in gene expression or cell fate, but also by rewiring the communication channels between pigment cells. From this perspective, pigment patterns become not only a readout of cell identity, but also a dynamic product of tissue-level signaling. This kind of redeployment likely contributed to the vast diversity of pigment patterns observed in teleosts and other vertebrates. That such mechanisms rely on electrical and chemical coupling via gap junctions, a system also fundamental in heart, brain or fins, highlights the deep evolutionary plasticity of cellular communication in shaping several traits.

The *Snowflake* mutation, by altering Gja5b function, impaired pigment cell communication and effectively reduced pigment cell diffusion. As supported by our modeling, this disruption affected short-ranged cell coordination and transformed sharp bar boundaries into irregular expanded regions. In anemonefish the white bar pattern remains stable in terms of shape during fish growth

– a behavior not expected from classical Turing mechanism, which scales with size, as observed in zebrafish. Recent analysis of pattern formation of teleost fishes suggested that the blending of patterns generated by Turing patterning can generate diversity (*4*) but our results suggest that another dimension should be added to this as positional cues or boundary information may complement self-organizing processes, as shown here in *A. ocellaris*. More broadly, our results showed that altering biophysical properties in the skin can generate strong pattern changes without modifying the pigment cell types or their gene expression. In other words, pattern diversity can emerge not only from which cell types are present, but also from how these cells are organized and interact within the tissue. This view is consistent with morphodynamic patterning models (*26*) where tissue-level dynamics and mechanical feedback shape pattern outcome. By modulating how information spreads between cells, gap junction networks could offer a versatile and evolvable means to shape pattern diversity. While their wiring and timing may vary between species, the underlying logic of using regulated cell-cell communication to organize pigment patterns seems largely conserved.

## Acknowledgments

We thank the OIST sequencing center as well as the imaging section. We are grateful for the support of our husbandry teams at OIST and Academia Sinica, as well as the Mutualized Aquariology Service at the Oceanological Observatory of Banyuls sur mer, France. We thank the High Throughput Genomics Core of the Biodiversity Research Center at Academia Sinica for performing the NGS experiments. The core facility is funded by Academia Sinica Core Facility and Innovative Instrument Project (AS-CFII-108-114). We specifically thank Dr. Mei-Yeh Lu, Mr. Kuan-Ying Li, and Ms. Pei-Lin Chao for their efforts and assistance. We thank Simon de Bernard and Laurent Buffat from Altrabio for the initial bioinformatic analysis of the genome resequencing data.

## Funding

National Institutes of Health grant R35 GM122471 (DMP)

JSPS Grant-in-Aid for Scientific Research C 24K09472 (MW)

JSPS Grant-in-Aid for Scientific Research B 22H02678 (MKi, VL)

OIST KICKS Program for Research Scholarship Fund (VL)

Agence Nationale de la Recherche, project SENSO ANR19-CE14-0010 (LB)

## Author contributions

Conceptualization: MKl, MKi, DMP, VL,

Data curation: MH, SDV, RR

Formal analysis: MH, SDV, EG, BMM, RR, SP, DMP

Methodology: RR, SP, MKi

Resources: LS-K, MKi

Investigation: MKl, SM, SHL, SDV, HWH, BMM, RR, MW, YL, LB, MKi, DMP

Methodology: RR, SP

Validation: MKi

Visualization: MK, RR, DMP

Funding acquisition: VL, DMP, MKi,

Supervision: DMP, MKi, VL

Writing – original draft: MKl, RR, SP, MW, MKi, DMP, VL

Writing – review & editing: MKl, EG, SDV, MW, LB, SP, MKi DMP, VL,

## Diversity, equity, ethics, and inclusion [optional]

### Competing interests

Authors declare that they have no competing interests.

### Data and materials availability

Raw reads of bulkRNAseq datasets are available to download under NCBI BioProject accession number PRJNA1168256. All images, as well as count matrices and codes are available at Zenodo **10.5281/zenodo.16892159**.

## Supplementary Materials

Materials and Methods

Supplementary Text

Figs. S1 to S7 Tables S1 to S3

References (*27*–*62*)

Movies S1 to S2

## Notes

### Competing Interest Statement

The authors have declared no competing interest.

