## Supplemental Material for "Cell-cell communication as underlying principle governing color pattern formation in fishes"

Supplementary Materials for “Cell-cell communication as underlying principle governing color pattern formation in fishes” include:

Materials and Methods
Supplementary Text
Figs. S1 to S7
Tables S1 to S3
References 27-62
Movies S1 to S2

### Materials and Methods

#### Terminology

In a cross with two *Snowflake* parents, the offspring showed Mendelian proportions of 25% wildtype, 50% heterozygous, and 25% homozygous mutants. All homozygous mutant larvae died shortly after hatching. Therefore, unless noted otherwise, throughout this paper wild-type (WT) refers to the phenotypically wild-type individuals coming from a *Snowflake* cross (25%) and *Snowflake* (SF) refers to the heterozygous individuals (50%).

Naming conventions between mammalian and zebrafish connexin genes and proteins are not congruent. In mammalians connexin genes are indicated by the prefix gj (for gap junction) followed by the class (e.g. a, b, c) and a number. The proteins, however, are named according to their differences in size using the prefix Cx followed by the predicted molecular weight. For example, the gene *gja5* corresponds to the protein Cx40. The naming system in zebrafish usually uses Cx followed by predicted molecular weight for both gene and protein. Unfortunately, the predicted molecular weights for mammalian and zebrafish connexin genes vary, which makes the naming systems confusing at times. Taking the example from above, mammalian protein Cx40 (gene *gja5*) is orthologous to zebrafish Cx41.8 (gene *Cx41.8*). To simplify, throughout this paper the prefix gj followed by the class is used for both gene (e.g. *gja5b*) and protein (e.g. Gja5b). The corresponding zebrafish Cx naming is only mentioned at the first reference.

#### Phenotypic comparison during development

To quantify white area expansion the wideness of the body bar was measured by counting the number of dorsal fin spines and dorsal fin soft rays the bar encompassed (Fig. 1A). Additionally, white areas were calculated as follows: First, using ImageJ an outline was drawn manually around the entire body (excluding fins) and the area was determined. Second, using ImageJ outlines were drawn manually around each of the three white bars (excluding fins) and the areas were determined (Fig. S7A). Lastly, the percentage of white areas in relation to the total body area were calculated either individually for each bar or combined (total white area). The same procedures were followed to calculate the orange and the black area, respectively.

The bending angles of the black edge were calculated using ImageJ (Fig. S7B). Wild-type *A. ocellaris* have one bending angle of about 120–150°, which is sometimes referred to as the "bulge". This is usually located on the anterior edge of the body bar, approximately on level with the midline.

The length of the black edge (body bar only) was calculated for both the anterior and posterior edge individually (Fig. S7C). First, a straight line was drawn to connect the dorsal and ventral points were the black met the body outline. Next, a curved line was traced along the actual black edge. Both line values were put into relations – with a value of 1 indicating the shortest possible outcome.

#### SNV analysis

The genomes of 20 wild-type (WT) and 20 heterozygous *Snowflake* (SF) sibling fish were sequenced. DNA extraction was done using Promega AS1400 Maxwell RSC Blood DNA Kit (Promega) following the manufacturer's instructions with an added RNase step. Library preparation (NEBNext® Ultra™ II FS DNA Library Prep Kit for Illumina) and DNA Shotgun Sequencing (150bp, pair ends) using NovaSeq 6000 S2 whole (NovaSeq 6000 S2 Reagent Kit v1.0, 300 cycles) was performed by the in-house sequencing center (OIST SQC, sample ID 137).

Raw genomic data were processed using the Genome Analysis Toolkit (GATK) framework (*27*) following Altrabio's (www.altrabio.com) recommended pipeline, a standardized analysis protocol that incorporates the necessary modifications for coping with the absence of reliable databases of known variants for this species. Briefly, quality of raw reads was assessed using FastQC v0.11.9 (https://www.bioinformatics.babraham.ac.uk/projects/fastqc/). Adapters, primers, low quality ends (Phred score < 30), and reads shorter than 35 bp were removed with Trimmomatic v0.36 (*28*) using the parameters ILLUMINACLIP:TruSeq3-PE.fa:2:30:10:2:true LEADING:3 TRAILING:3 SLIDINGWINDOW:4:15 MINLEN:36. Sample contamination was evaluated using FastQ Screen v0.14.0 (https://www.bioinformatics.babraham.ac.uk/projects/fastq_screen/). The *A. ocellaris* reference genome (*29*) was indexed using Bowtie2 v2.4.4 (*30*) and reads were aligned using the bwa-mem2 v2.2.1 (*31*) algorithm. The resulting alignments were sorted, marked for duplicates, and indexed using Picard tools v2.22.1 (https://github.com/broadinstitute/picard). Next, the GATK HaplotypeCaller tool was used to call variants per-sample, including summarized evidence for non-variant sites (GVCF workflow). The variant-calling matrices of the different alignments were then imported into GenomicsDB using GATK GenomicsDBImport and all samples were jointly genotyped with GATK GenotypeGVCFs. Variants were filtered based on the following criteria: " QD2" = QD < 2.0, "SOR3" = SOR > 3.0, "FS60" = FS > 60.0, "MQ40" = MQ < 40.0, "MQRS125" = MQRankSum < -12.5, "RPRS8" = ReadPosRankSum < -8.0 for SNVs, and "QD2" = QD < 2.0, "SOR3" = SOR > 3.0, "FS200" = FS > 200.0, "RPRS20" = ReadPosRankSum < -20.0 for indels. This resulted in a raw genotype matrix counting 5,189,387 SNVs and 1,416,665 indels (83.8% and 90.5% of the SNVs and indels that passed filtering, respectively). Subsequently, to annotate the variants, the reference genome (*29*) was annotated using the *A. ocellaris* genome and annotations from Ensembl. Mapping of the coordinates between the two assemblies and transfer of the Ensembl annotations onto the reference assembly were performed using Liftoff v1.6.1 (*32*) with the option -polish. The software SnpEff v5.0 (*33*) was then used to predict the putative effect of all variants. Finally, after annotating the variants, these were further filtered by counting the number of heterozygous SF and homozygous WT at each position. The sum of these values generated a "heterozygosity score" between 0 (all-homozygous SF and all-heterozygous WT) and 40 (desired pattern of all-heterozygous SF and all-homozygous WT). Only variants in coding regions and with a score greater than 35 were considered further.

SNV confirmation

Sanger sequencing was employed to rule out genes that are not primarily causing the *Snowflake* phenotype. Fish from three *Snowflake* strains, originating either in France, Taiwan, or Japan, were fin clipped. DNA was extracted employing the Promega AS1400 Maxwell RSC Blood DNA Kit (Promega) following the manufacturer's instructions with an added RNase step. PCRs were performed for each of the seven candidate genes (primers details in Table S3) using either KOD FX (TOYOBO) or Go Taq (Promega) PCR kits. PCR conditions for KOD FX were the following: first denaturation (94°C for 2 min), 30 cycles of annealing and elongation (98°C for 10 sec, 58°C for 15 sec, 68°C for 40 sec), and a final elongation (68°C for 5 min). PCR conditions for Go Taq were the following: first denaturation (95°C for 2 min), 30 cycles of annealing and elongation (95°C for 30 sec, 65°C for 30 sec, 72°C for 40 sec), and a final elongation (72°C for 5 min). PCR products were purified using Exo-CIP™ Rapid PCR Cleanup (NEB). An automated Capillary sequencer (Thermo Fisher 3500xl Capillary sequencer) in combination with the primers listed above for the seven candidate genes and the BigDye Terminator v3.1 Cycle Sequencing Kit (Thermo Fisher) were used for direct Sanger sequencing.

For each gene between 10-20 wild-type and 10-20 *Snowflake* fish were tested (Table S1). A gene could be excluded from the *Snowflake* causing gene list, if either one of the following were true: (1) a phenotypic *Snowflake* offspring exhibited a homozygous wildtype genotype for a specific gene, or (2) a phenotypic wildtype offspring had a heterozygous mutant genotype for a specific gene (and a homozygous mutant genotype for this gene was never observed).

<u>Phylogenetic Analysis</u>

Sequences for all available *gj* genes for *Danio rerio* and *Amphiprion ocellaris* were downloaded from NCBI. Sequences for *gja5* were downloaded for three additional species: *Oryzias latipes*, *Gasterosteus aculeatus*, and *Takifugu rubripes*. Conserved regions for *Danio rerio* were acquired from Mikalsen and colleagues 2020 (*34*) and all individual NCBI sequences were aligned and trimmed to those conserved regions in SeaView (PRABI-Doua: SeaView). A NJ phylogenetic tree was created using MAFFT v7 (https://mafft.cbrc.jp/alignment/server/index.html) selecting JTT substitution model and Bootstrap resampling of 1000.

<u>Pharmacological experiments</u>

Five to eight pre-metamorphic larvae between 6-8dph (days post hatching) were placed into a 500mL glass beaker (as described in (*35*)) and incubated with commercially available pharmacological active small molecules. The following small molecules were used in this study: 0.3-0.6mM putrescine (putrescine dihydrochloride P5780, Sigma-Aldrich), 0.2-0.6mM spermidine (spermidine S0266, Sigma-Adrich), 0.3-0.6mM spermine (spermine S4264, Sigma-Aldrich) and 50-300nM PQ7 (gap junction enhancer PQ7 5.06216, Sigma-Aldrich). The larvae were raised following previously published protocols (*35*). Every day 20% of water was changed and replaced with seawater including the corresponding small molecule in the desired concentration. Incubation time varied between 5-10 days. To end the experiment the concentration of the small molecule was dramatically lowered. First, 80% of the drug infused water was replaced with normal sea water, adding the water in small increments over several hours. Second, the following day 50% of the water was replaced with normal sea water. Imaging of larvae started the next day, three days after the end of the experiment. The larvae were anesthetized with 100mg/L MS222 (Sigma-Aldrich), placed in a petri dish and images of both sides were taken using a Discovery.V20 stereoscope (Zeiss) mounted with a Axiocam 208 color camera (Zeiss). Images of the same fish were taken successively, every 2-5 days until all three bars were formed.

<u>CRISPR/Cas9 mutagenesis</u>

For amino acid substitution, homologous gene knock-in method using CRISPR/Cas9-nickase system was employed. For primer details see Table S3.

**Construction of SF- plasmid donor** (Fig. S2E): A 697 bp fragment-SF, which contained *gja5b* exon 4 including the E42K mutation and its flanking regions, was amplified by PCR. A fragment-L was amplified by the primers Ao-gja5-HAL-XhoI-F and Ao-gja5-SF-R with genomic DNA as template. A fragment-R was amplified by the primers Ao-gja5-mut-F and Ao-gja5-HAR-SpeI-R with genomic DNA as template. Then, a mixture of fragments L and R was used as template to amplify fragment-SF by PCR with primers Ao-gja5-HAL-XhoI-F and Ao-gja5-HAR-SpeI-R. As a result, fragment-SF contained an amino acid substitution from Glu (GAG) to Lys (AAA) and 11 silent mutations in the coding region (small orange letters in Fig. S7D), which enabled the designated design of specific primers for the detection of gene substitution ("Ao-KI-check-R" in Table S3). The PCR mixture contained 1μL of each primer (2μM) and 0.2μL of KOD-FX DNA polymerase (TOYOBO) in a total volume of 10μL. The PCR conditions were as follows: 94°C for 2 min followed by 30 cycles of 98°C for 10 sec, 55°C for 10 sec, and 68°C for 30 sec. The fragment-SF and p2BaitD-acta1_500 bp-mAG (*36*) were digested with Xho I and Spe I and ligated to generate SF-donor plasmid. The donor plasmid contained two invert-repeat of BaitD sequence with which the knock-in fragment is cut out by Cas9-nickase (*36*).

To confirm successful substitution, skin from a G0 individual was collected, with one sample taken from a normal white bar (CRISPR fish WT bar) and another sample taken from a *Snowflake* reminiscent white bar (CRISPR fish SF bar). DNA from the skin tissues was extracted using Proteinase K and phenol/Chloroform. The substituted sequence was confirmed by PCR and followed by Next-generation sequencing. PCR was performed with the forward primer Ao-gja5-KI-check-F2, which is located outside of the fragment-SF, and the reverse primer Ao-KI-check-R, which was designated to target the silent mutation sites (Table S3). The amplicon was sequenced with NextSeq 1000 and analyzed subsequently. The reference *Amphiprion ocellaris* genome assembly ASM2253959v1 ((*29*); as downloaded from NCBI) was indexed with the "build" function of bowtie2 v2.4.4 (RRID:SCR_016368; (*30*)). Raw sequencing reads were then aligned to this genome, using bowtie2 with default parameters. The resulting .sam files were sorted and converted to .bam files, using functions "sort" and "fixmate" from SAMtools v1.18 (RRID:SCR_002105; (*37*)) with default parameters. The resulting files could be imported into the Integrative Genomics Viewer (IGV) Desktop Application (RRID:SCR_011793; (*38, 39*)), therefore obtaining pileup plots. Base percentages at each position of the *gja5b* coding sequence (LOC111585045) were obtained directly from the application and visualized as an Excel table. The introduced donor plasmid will induce a replacement of the codon 42 from GAG to AAA, if successful. Sequencing results showed a 0% substitution rate for wild-type fish, while the CRISPR G0 fish displayed some level of substitution: 1% for the WT bar and 6% for the SF-bar (Fig. S2F). The rate of the silent mutations in the CRISPR fish SF-bar also reached 6%, suggesting this was the percentage of successfully achieved genomic edits.

**Microinjection**: A mixture of Ao-gja5-crRNA2 (40ng/μl, Target seq: TACGGCAGCCGAGTCTTCGTGGG, PAM), BaitD-crRNA (40ng/μl, Target seq: GATCTTCGGCCTAGACTGCGAGG), tracrRNA (140ng/μl, FASMAC), Cas9-nickase protein (D10A)(1,000ng/μl, FASMAC), and the donor plasmid (2.5ng/μl) was microinjected into the cytoplasm of 1- or 2-cell stage fertilized eggs as described previously (*40*). Eggs were raised in egg tumblers and then transferred to tanks. Fin clips were cut after 2-3 months and the presence of induced mutations within *gja5b* was checked via PCR (see above).

**Random *gja5b* mutagenesis:** For the induction of random mutations in *gja5b*, a single guide RNA (sgRNA2), targeted the same sequence as Ao-gja5-crRNA2, was synthesized with CUGA gRNA synthesis kit (NIPPON gene) following the manufacturer's instructions. Then, the sgRNA2 (100ng/μl) and Cas9 protein (500ng/μl, FASMAC) were microinjected into 1-cell stage of fertilized eggs as described previously (*40*). To confirm successful mutagenesis, five scales from white bars showing a *Snowflake* phenotype were picked with forceps from five CRISPR/Cas9 G0 fish. As a control, five scales were picked from wild-type white bar coming from one non-treated fish. Scales were incubated in 8μl of 25mM NaOH and 0.5mM EDTA at 95°C for 10 min. After neutralization by addition of 10μl 40mM Tris-HCl, pH8.0, the resulting solutions were used as template for PCR. PCR was performed using KOD-FX DNA polymerase (TOYOBO) with primers, TruSeq-F and TrySeq-R (Table S3), both contain adaptor sequence for TruSeq sequencing. PCR products were purified with NucleoSpin Gel and PCR Clean-up (Macherey-

Nagel) following instructions. Then, sequencing was performed with NextSeq 1000. From each individual, more than 32000 amplicon reads were obtained. In the white bar of the wild-type fish, no mutated reads were observed, only wild-type sequences. On the contrary, white bar areas of the five CRISPR treated fish contained many types of mutations, including short in-frame deletions (less than 19 nucleotides). Those were particularly frequent representing 22 – 52% of all reads in these fish (Fig. S2G) and believed to mainly contribute to the *Snowflake*-like phenotype.

Visualization of *gja5b* expression pattern

*gja5b* expression pattern was visualized with green fluorescence using Tol2 transposon system. A GFP expression plasmid driven by *gja5b* promoter, pTol2-Aogja5bPro6KGFP (Fig. S3B) was generated. The upstream region (ca. 6.6kb) from translation codon (ATG) of *gja5b* was amplified by PCR using PrimeStar GXL DNA polymerase (Takara Bio) and primers Ao-gja5bPro-F0 and Ao-gja5bPro-R1 (Table S3). The Tol2 trap vector, pT2AL200R150 (*41*), was digested with Xho I and BamHI to remove intrinsic sequence of EF1a-P and the intron. The PCR fragment and digestive of pT2AL200R150 were ligated using In-Fusion HD Cloning Kit (Takara Bio). Tol2 transposase mRNA was synthesized from pCS-TP (*42*) using an mMessage mMachine SP6 Kit (Ambion). The mixture of pTol2-Aogja5bPro6KGFP (5ng/μl) and Tol2mRNA (50ng/μl) was microinjected into fertilized wild-type eggs (see above).
GFP fluorescence was observed using a fluorescence stereomicroscope (SZX16, OLYMPUS) with GFP filter set (excitation: BP460-495, emission: BA510IF).

Stage-wise transcriptomics

**RNA extraction and sequencing:** 10-15 Larvae of all seven identified postembryonic stages (*43*) were euthanized in 100mg/L MS222, imaged with an Axiocam 208 color (Zeiss) mounted on a Discovery.V20 stereoscope (Zeiss) and stored in RNAlater (Sigma-Aldrich). RNA was extracted using AS1340 Maxwell RSC simplyRNA Tissue Kit (Promega) following the manufactures instructions. Library preparation (NEBNext® Ultra™ II Directional RNA Library Prep Kit for Illumina) and RNA Sequencing with poly-A RNA Purification (strand-specific, 150bp, pair ends) using NovaSeq 6000 S2 whole (NovaSeq 6000 S2 Reagent Kit v1.5, 300 cycles) was performed by the in-house sequencing center (OIST SQC, sample ID 382).

**Analysis**: The complete Rnotebooks use for the analysis of the stage-wise transcriptomic data (also indicating the versions of all packages used), the counts matrix, the gene and sample metadata files, as well as other data to reproduce the analysis are available at Zenodo **10.5281/zenodo.16892159**; the pipeline is summarized below. Raw (demultiplexed) fastq files were quality-checked based on reports generated by using FastQC v0.11.9 (RRID:SCR_014583; https://www.bioinformatics.babraham.ac.uk/projects/fastqc/) with default parameters, before and after adapter trimming. Adapters were trimmed using Trimmomatic v0.39 (*28*) with parameters ILLUMINACLIP:TruSeq3-PE-2.fa:2:30:10:8:keepBothReads LEADING:3 TRAILING:3 SLIDINGWINDOW:4:15 MINLEN:36. Trimmed reads were quantified at the transcript-level using the pseudo-aligner kallisto v0.46.2 (*44*) against the *A. ocellaris* reference transcriptome. The average mapping rate was 64.03% of total reads. The counts data was processed as a DGEobject (edgeR package, RRID:SCR_012802; (*45-47*)) and differences in library size were taken into account by obtaining counts per million (CPM) values (edgeR's function cpm) to allow a comparable threshold to filter out of lowly expressed genes. Here, we only maintained genes for which at least 10 counts could be detected (arbitrary) in at least 10 samples (our smallest experimental unit being n=10 per treatment). This corresponded to a threshold of 0.95 CPM based on the sample with lowest library size, resulting in a filtering out of 1755 out of 14949 genes (11.7%). Heatmap visualizations were computed on the filtered count data, taking into account library size differences and correcting against compositional bias through the Median of ratios method internal to the DeSeq2 pipeline (DESeq2 package, RRID:SCR_015687; (*48*)). Counts were then further variance stabilized (DESeq2's varianceStabilizingTransformation function). All heatmaps were plotted using the function Heatmap from the ComplexHeatmap package (*49, 50*) reporting z-scores ((counts − mean)/sample standard deviation; base R's "scale" function with default parameters).

<u>Scale transcriptomics</u>

Scales (around 100) of each color were manually collected using forceps and transferred into sterile 2mL microcentrifuge tubes filled with 750mL ice-cold Trizol (TRI Reagent; Merck/Sigma-Aldrich) containing three autoclaved stainless steel beads (EBL Biotechnology). Samples were disrupted and homogenized by mechanical agitation in a vibrating bead mill (TissueLyser II, Qiagen, RRID:SCR_018623; 3 min, 30Hz, room temperature). Homogenized sample lysates were stored at -80°C before RNA extraction. On the day of extraction, sample lysates were thawed on ice and centrifuged 10 min in a tabletop microcentrifuge at 11000 x g, 4°C; as to collect any leftover tissue debris to the bottom. 750µL of each supernatant were loaded into a dedicated RNA filtering column (NucleoSpin® RNA Mini kit; Macherey-Nagel) and processed according to the manufacturer's recommendations. Specifically (for each sample) 350mL of 70% ethanol (Honeywell/Riedel-deHaen) were added to the filtered flowthrough, the solution was then vortexed, and loaded onto an RNA binding column. After desalting (kit-supplied Membrane Desalting Buffer), contaminating DNA was digested by a 15 min incubation in recombinant DNAse (in Reaction Buffer, room temperature). Guanidine hydrochloride/ethanol wash buffers (RAW2 and then, twice, RA3) were then sequentially used to denature proteins, discarding the flowthrough from centrifugation after each step. After a further centrifugation to remove all possible leftover buffer/ethanol (2 min, 11000 x g, 4°C), membrane-bound RNA was eluted in 30mL of RNAse-free water (Invitrogen™/Thermo Fisher Scientific), in a 1.5mL RNAse-free tube (kit-supplied). RNA amounts and quality were initially assessed with a NanoDrop spectrophotometer (Thermo Scientific™) and by agarose gel electrophoresis (2:1 intensity ratio of 28S:18S bands; 800ng of sample per lane, 0.6 % Agarose in TAE-buffer, 1h, 50V). Extracted RNA was stored at -80C until sequencing.

Quality-control, library preparation and sequencing of extracted scale and skin RNA were performed by the High Throughput Genomics Core at Academia Sinica, Taipei. RNA-Seq libraries were generated from 1000ng of total RNA (or lower for few samples for which this amount could not be reached) using the Illumina Stranded mRNA Prep mRNA Sample Preparation Kit with UDI indices (Illumina) according to manufacturer's instructions. Surplus PCR primers were removed using AMPure XP Bead-Based Reagent (Beckman Coulter Life Sciences). Final cDNA libraries were checked for quality and quantified using Qubit (ThermoFisher Scientific) and Fragment Analyzer for size profiling (Agilent), and concentration-normalized using KAPA Library Quantification Kit for Illumina Platforms (Roche). Sequencing was performed on an Illumina NextSeq2000 for paired-end 150 base format. Libraries were loaded in a P3 flow cell. The fastQ files were generated and demultiplexed using the Illumina bcl2fastq v2.20 pipeline. A median of 29 million paired-end reads per sample and 26 million paired-end reads per sample were obtained (expected output ceiling: 31 million reads/sample and 27 million reads/sample, respectively). Three samples, which had shown low qPCR amplification rates during library preparation, produced a lower data yield than other samples, despite equal pooling. A further sequencing round was thus run for these samples (on a P2 flowcell), producing an average of 15 million extra reads for each. The complete Rnotebooks use for the analysis of the scale and skin transcriptomic data (also indicating the versions of all packages used), the counts matrix, the gene and sample metadata files, as well as other data to reproduce the analysis are available at Zenodo **10.5281/zenodo.16892159**; the pipeline is summarized below.

**Pre-processing**: Raw (demultiplexed) fastq files were quality-checked based on reports generated by using FastQC v0.12.0 (RRID:SCR_014583; (*51*)) with default parameters, before and after adapter trimming. Adapters were trimmed using the function bbduk (RRID:SCR_016969) of BBTools v39.01 (RRID:SCR_016968; Bushnell B., http://sourceforge.net/projects/bbmap/) with ktrim=r, and k=23, qtrim=r, trimq=30, trimpolyg=40. Categories flagged by FastQC after trimming ("warning" or "fail") were analyzed in detail and judged not to be prejudicial to further analysis.

**Analysis**: For scale transcriptomic data, trimmed reads were quantified at the transcript-level using the pseudo-aligner salmon v1.10.1 (RRID:SCR_017036; (*52*)) against the *A. ocellaris* reference transcriptome, using the genome as a decoy (decoy-aware pseudo-alignment; assembly ASM2253959v1; (*29*)), as per documentation, and using flags --validateMappings -- seqBias -- gcBias. The average mapping rate was above 86% of total reads. Salmon output transcript-based quantification files (quant.sf) were imported in RStudio (RRID:SCR_000432; Posit team, 2023) and summarized at the gene level using the tximport function from the tximport package (RRID:SCR_016752; (*53*)), referencing the gene models of the *A. ocellaris* reference genome assembly ASM2253959v1. The counts matrix was obtained by re-calculating counts through the flag countsFromAbundance = "lengthScaledTPM". Gene metadata (names, descriptions) were loaded from a custom-curated reference file based on the integration of Ensembl (still based on assembly AmpOce1.0) and NCBI annotations. This metadata reference is available at the code repository associated with this publication. Counts were processed as a DGEobject (edgeR package, RRID:SCR_012802; (*45-47*)) and differences in library size were taken into account by obtaining counts per million (CPM) values (edgeR's function cpm) to allow a comparable threshold to filter out of lowly expressed genes. Here, we only maintained genes for which at least 10 counts could be detected (arbitrary) in at least 2 or 3 samples. For the Okinawa dataset, this corresponded to a threshold of 0.46 CPM based on the sample with lowest library size, resulting in a filtering out of 4961 out of 26889 genes (18.4%). For the Taiwan dataset, this corresponded to a threshold of 0.51 CPM based on the sample with lowest library size, resulting in a filtering out of 6511 out of 26889 genes (24.2%).

Heatmap visualizations were computed on the filtered count data, considering library size differences and correcting against compositional bias through the Median of ratios method internal to the DeSeq2 pipeline (DESeq2 package, RRID:SCR_015687; (*48*)). Counts were then further variance stabilized (DESeq2's varianceStabilizingTransformation function). Finally, count data was corrected for latent sources of variation not related to pigment type, tissue of origin, or anemonefish strain, using the removeBatchEffect function from the limma package (RRID:SCR_010943; (*54*)) and as identified through the svaseq function from the sva package (RRID:SCR_012836; (*55*)). All heatmaps were plotted using the function Heatmap from the ComplexHeatmap package (*49, 50*) reporting z-scores ((counts − mean)/sample standard deviation; base R's "scale" function with default parameters).

Xenopus oocyte experiments

**Plasmid construction:** Tissue of juvenile fish was homogenized by mechanical agitation in a vibrating bead mill (TissueLyser II, Qiagen; 3 min, 30Hz, room temperature). Homogenized sample lysates were stored at -80°C overnight for RNA extraction. Total RNA was extracted using TRIzol Reagent Trizol (Merck/Sigma-Aldrich). Sample RNA and d(T)23VN (ProtoScript® II First Strand cDNA Synthesis Kit, NEB) were denatured for 5 mins at 65°C, spun in briefly and kept on ice. Manufacture's recommendations for ProtoScript II were followed to synthesize cDNA, which was used for subsequent PCR. PCR was performed using a Biometra Tone, with 10X KOD Plus Neo Buffer, 2mM each dNTPs, 25mM MgSO4, KOD-Plus-Neo Polymerase, 100ng template, and 50Um primer sets to a total volume of 50µl. The reaction conditions were as follows: 95°C for 5 mins; 35 cycles of 95°C for 30 sec, 60°C for 45 sec, and 72°C for 1 min; and final extension at 72°C for 10 mins. DNA fragments were electrophoresed on agarose gels, before being purified and ligated into the Zero Blunt TOPO PCR Cloning vector (Invitrogen). The resulting plasmids were used to transform *Escherichia coli* DH5$\alpha$ following standard protocols.

**cRNA preparation:** The cDNA fragments of *A. ocellaris* connexins were amplified with KOD-FX polymerase (TOYOBO). Primer details can be found in Table S3. The amplified fragments were digested with EcoRI and NotI, and ligated into the pGEM-HeFx plasmid. The resulting plasmids were linearized using appropriate restriction enzymes and then used as templates for *in vitro* cRNA synthesis with mMESSAGE mMACHINE T7 Transcription Kit (Invitrogen), according to the manufacturer's protocol.

For the zebrafish *cx41.8* E42K mutant, the G124A DNA substitution was introduced into the previously used pGEM-HeFx-zfcx41.8 plasmid (*15*) by PCR with KOD-Fx polymerase. The primer sequences are found in Table S3. The resulting PCR product was used for transformation of *E. coli*. After selection, plasmids were isolated from colonies, and those containing the confirmed G124A DNA substitution were used as templates for *in vitro* cRNA synthesis.

**Preparation of *Xenopus* oocytes:** Whole cell voltage clamp recording with *Xenopus* oocytes were performed as described (*15*). An adult *Xenopus laevis* female was anesthetized with MS222 (Sigma-Aldrich), and the ovarian lobes were collected using surgical knife and forceps. The eggs were treated with collagenase solution (20mg/ml collagenase I (Sigma) and 20mg/ml hyaluronidase (Sigma) in OR2 buffer (82.5mM NaCl, 2mM KCl, 1mM MgCl2, and 5mM HEPES [pH 7.5, adjusted with NaOH])) at 18°C for two hours. Stage V and VI oocytes were collected manually and used for cRNA injection. 0.1-10ng of cRNA was co-injected with 10ng of antisense oligonucleotide DNA for *Xenopus* Cx38 into *Xenopus* oocytes. Water was co-injected with the antisense oligonucleotide as a negative control. Oocytes injected with cRNA were incubated at 18°C overnight in ND96 buffer (93.5mM NaCl, 2mM KCl, 1.8mM CaCl2, 2mM MgCl2, and 5mM HEPES [adjusted to pH 7.5 using NaOH]). The vitelline membrane was removed manually using forceps in a hypertonic solution (200mM aspartic acid, 1mM MgCl2, 10mM EGTA, 20mM KCl, and 10mM HEPES [pH 7.5]) and the oocytes were manually paired with the vegetal poles together in the handmade agarose chamber and incubate at 18°C overnight in ND96 buffer. For electrophysiological analysis of heterotypic gap junctions, oocytes injected with different connexin cRNA were paired after removal of the vitelline membrane.

**Transjunctional (gap junction) current recording:** Transjunctional currents were measured using the dual whole-cell voltage clamp technique with two iTEV90 multielectrode clamp amplifiers. Current and voltage electrodes were prepared with a micropipette puller P-1000 (Sutter Instrument) to obtain a resistance of 0.5–1.0MΩ. The pipette was filled with solution containing 3M KCl, 10mM EGTA, and 10mM HEPES (pH 7.4). Both cells were initially clamped at −40mV and one cell was then subjected to 3-s voltage steps from −140 to +60 mV in 20mV increments. Currents detected in the second oocyte were recorded and the current values at the end of the steady state of -100mV were compared.

Zebrafish experiments

**Fish stocks:** The semi-dominant zebrafish allele *gja5b*$^{stl710}$ was obtained in an ethynitrosorurea F3 forward genetic screen (*56*) and maintained subsequently in the AB* background. The mutant was mapped by association with variants in identified by whole-genome resequencing of pooled genomic DNA of wild-type and mutant individuals (*20, 57*), with confirmation of mutant lesion by Sanger sequencing. Subsequent phenotypic analyses of *gja5b*$^{stl710}$ used fish maintained in AB*.

**Plasmid construction:** *defbl1:gja5b*$^{WT}$*-DrIRES-nEos*, *defbl1:gja5b*$^{stl710}$*-DrIRES-nEos*, *mitfa:gja5b*$^{WT}$*-DrIRES-nEos* and *mitfa:gja5b*$^{stl710}$*-DrIRES-nEos* plasmids were assembled using Gibson Assembly Master Mix (NEB E2611) according to manufacturer's protocol. Plasmids containing regulatory sequences of *mitfa* (melanophore and xanthophore lineage) and *defbl1* (iridophore and melanoleucophore lineages) have been described (*19, 20*). Constructs used destination plasmid Tol2pA2 from the Tol2kit as backbone (*58*); coding sequences of *gja5b*$^{wt}$ or *gja5b*$^{stl710}$ were cloned from genomic DNA.

**Microinjection:** Plasmids were injected at one-cell stage with *tol2* mRNA at 25 ng per embryo. Injected embryos were reared in accordance with the recommendations in the Guide for the Care and Use of Laboratory Animals of the National Institutes of Health and approved institutional Animal Care and Use Committee (ACUC) protocol (#4170) of the University of Virginia.

**Imaging and documentation:** Fish were anesthetized in MS222 (Syndel USA) prior to imaging. In some microscopic images, fish older than 5dpf were anesthetized and treated with 1mg/mL epinephrine (Sigma-Aldrich) to contract pigment granules. Whole adult fish images were acquired using a Nikon D810 full-frame DSLR camera with a 105 mm MicroNikkor macro lens. All other bright-field images were collected with Zeiss AxioZoom stereomicroscope, or Zeiss AxioObserver inverted microscope equipped with Zeiss Axiocam color cameras using ZEN Blue software. For developmental time series of *gja5b*$^{stl710}$ mutants and wild-type siblings, fish were imaged daily though stages of adult pattern formation as described (*59, 60*), with melanin contracted to cell centers by pre-treatment with epinephrine. For easier visualization of lightly melanized cells in schematics, brightfield color images were binarized and pseudocolored and boundaries of individual melanin spots representing individual cells were enlarged uniformly by processing in Adobe Photoshop. Other images were color balanced and adjusted for display levels in Adobe Photoshop. Quantification of pattern variation in transgenic zebrafish fish followed the same methods as used for anemonefish, described above.

**Gap junction gene expression in zebrafish:** Re-analyses of single nuclear gene expression data from (*61*) for dot-plot of gap gene transcript abundance in pigment cell and other cells during adult pattern formation used built-in functions of Monocle 3 (*62*).

### Edwards-Wilkinson modeling

**Bar profile extraction and preprocessing:** From each image, the dorsal-ventral bar was segmented. For each row $i = 1,\ldots, R$ of the grayscale mask were located the leading-edge coordinate $x_i$ and computed the local height

$$h_i = \frac{1}{\sum_j I_{i,j}} \sum_j j I_{i,j} \times \Delta y,$$

where $I_{i,j}$ is the grayscale intensity at pixel $(i, j)$ and $\Delta y$ is the calibrated pixel height. Each profile was then rotated and vertically shifted so that $h(0) = h(L) = 0$, where L is the total physical bar length.

**Height-difference variance:** For each profile a band pass filter was applied to isolate the modes in which the power spectrum followed $k^{-2}$. The filtered spectrum was transformed back into real space and the height-difference variance was computed

$$V(\Delta x) = \langle [h(x + \Delta x) - h(x)]^2 \rangle_x.$$

**Analytical prediction and estimation of $\sigma^2/D$:** The one-dimensional Edwards-Wilkinson (EW) model predicts the exact height-difference variance is

$$V_{int}(\Delta x) = \frac{2}{\pi} \frac{\sigma^2}{D} \int_{k_{min}}^{k_{max}} \frac{1 - cos(k\Delta x)}{k^2} dk,$$

where $\sigma^2/D$ is the noise-to-surface-tension ratio. To estimate $\sigma^2/D$ from data, the power spectrum was computed (Fig. S6A-D)

$$P(k) = \frac{1}{N} |H(k)|^2, H(k) = \sum_{j=1}^{N} h(x_j) e^{-ikx_j},$$

and fitted in the band $k_{min} \le k \le k_{max}$ to the scaling law

$$P(k) \sim \frac{\sigma^2}{D} \frac{1}{k^2}.$$

From this an empirical value of $\sigma^2/D$ was obtained.

**Statistical analysis:** All variance curves were computed for each of $n = 10$. Means and standard errors are reported.

### Supplementary Text

Snowflake mutants are defective for pattern boundary positioning

The *Snowflake* mutation of *A. ocellaris* is assumed to have appeared only once in aquaculture (1999 Tropical Marine Centre, UK) and then spread in the pet shop trade. A *Snowflake* pair produces about two-thirds *Snowflake* offspring and one-third wild-type offspring, suggesting a dominant mutation having an early lethal phenotype when homozygous.

Expansion of white bar areas was evident dorsally, where the numbers of white spines and soft rays of the dorsal fin were significantly increased in two-month juvenile *Snowflake* fish compared to wild-type fish (Fig. S1E). Total areas covered by white bars were also significantly increased, comprising ~50% of the total body in *Snowflake* as compared to only ~30% in wild-type (Fig. 1C). In *Snowflake* individuals the black boundary between orange and white areas is uneven and jagged. The black edge in wild-type *A. ocellaris* is relatively straight with no bends under 150°, except for the anterior bulge on the body bar (circled area in Fig. 1A) which is on average 149 degrees (± 5.6°). On the contrary, the black edge of *Snowflake* juveniles possesses several bends between 120-150° (Fig. S1F). Moreover, even bends between 70-120° are readily observed in *Snowflakes* (Fig. S1F), but never in wild-type. Another way to assess the unevenness of the black edge is to calculate the total distance from dorsal to ventral edges (see methods). In *Snowflakes* the black edge is considerably longer (Fig. S1G) than in wild-type, due to the jaggedness. The *Snowflakes'* jaggedness of the black edge becomes more pronounced as the color pattern matures (Fig. 1D).

Black edges in *Snowflake* are also thicker. The dominating body color of wild-type juveniles (2 months) is orange (on average 63%), followed by white (on average 30%) and black (on average 7%) (Fig. 1C). By contrast, *Snowflake* juveniles have nearly twice the black body area (on average 13%). This increased black area mainly results from increased thickness of the edges.

A connexin gene, *gja5b*, is mutated in *Snowflake*

To identify the gene (or genes) underlying the *Snowflake* phenotype, we used next generation sequencing of whole genomes from 20 phenotypically wild-type and 20 phenotypically *Snowflake* siblings derived from a backcross using the same wild-type female (Fig. S2A). Single nucleotide variants (SNVs) were found in 5 of the 24 chromosomes, chromosomes 2, 16, 17, 20, and 22. The largest cluster of SNVs was located towards the end of chromosome 16, including 1828 variants with the maximal heterozygosity score (see material & methods). We focused on the region of chromosome 16 that stood out with the largest number of high scoring SNVs (Table S2 shows the location/type of variant for the 1828 variants). Missense variants were deemed to be highly likely to affect RNA and/or protein features in mutants and were therefore scrutinized in further detail.

Cell-cell communication governs color pattern formation in anemonefish

Transcriptomic data sets from color-specific scale and skin samples were analyzed. Both data sets showed similar results, but skin samples, especially black skin, comprised other pigment cells and were less pure for single chromatophores. Therefore, we focused our analysis mainly on transcriptomic data acquired from scales. Because the purity of black color was limiting, we added samples from the *Black A. ocellaris* mutant strain in which the orange body color is replaced with black.

Even though most of the *gja5b* expression domains are evident in the white bar areas, we found minor reporter expression sites in orange skin (74%) (Fig S3C), which is populated by xanthophores and melanophores. Additional minor expressions were detected in fin rays (53%) (Fig S3C), lens (47%), vasculature (32%) (Fig S3C) and scales (32%). Therefore, we cannot formally exclude broader expression, and we do not claim iridophore-specific expression.

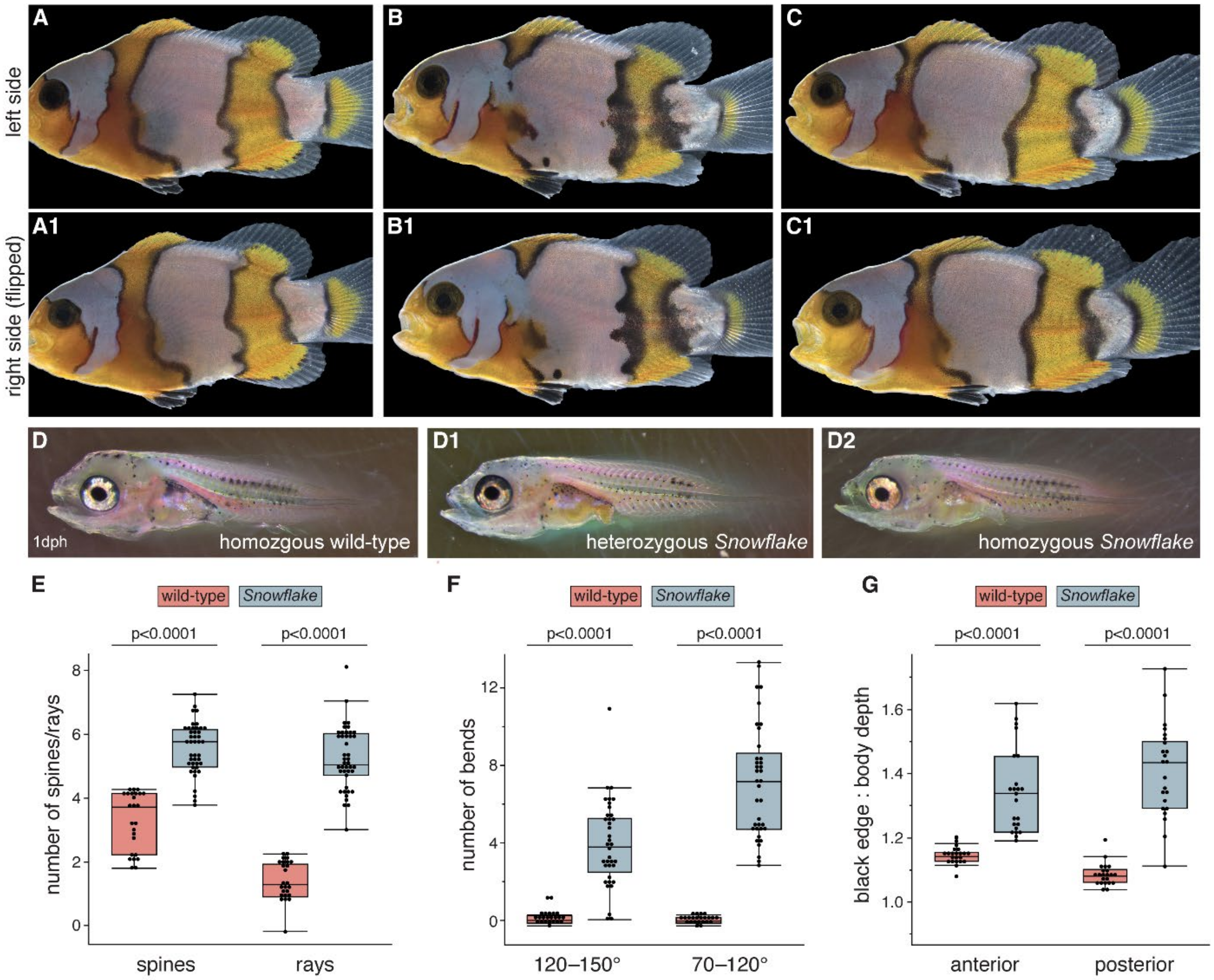

**Fig. S1. Features of the *Snowflake* phenotype.**
(**A** to **C**) Even though the *Snowflake* pattern varies considerably between individuals (A to C), the pattern of a single fish is highly bilateral symmetrical (compare top and bottom rows). (**D**) *Snowflake* larval color patterning (before metamorphosis) is indistinguishable from wild-type fish at 1 day post-hatching (dph). (**E**) The number of dorsal fin spines and rays included in the white area of the trunk bar is significantly greater in *Snowflake* fish (blue-grey bars) than in wild-type (pink bars), reflecting the greater anterior-posterior width of this bar in mutant. (**F**) The trunk bar in wild-type fish exhibits a single bend of 140–150°, the so-called anterior bulge. By contrast, *Snowflake* fish exhibit several bends of 120–150° and 70–120°, reflecting the jagged outline of their bars. (**G**) Another measurement of the *Snowflake's* jagged bar outline is the length of the black edge relative to body depth (a measure of overall body size); this ratio is consistently greater in *Snowflake* than in the wild-type. Kruskal-Wallis tests of significance in E–G; meristic counts in E and F are jittered within ±0.3 of original values by uniform random distribution to facilitate visualization of individual data points, representing individual fish.

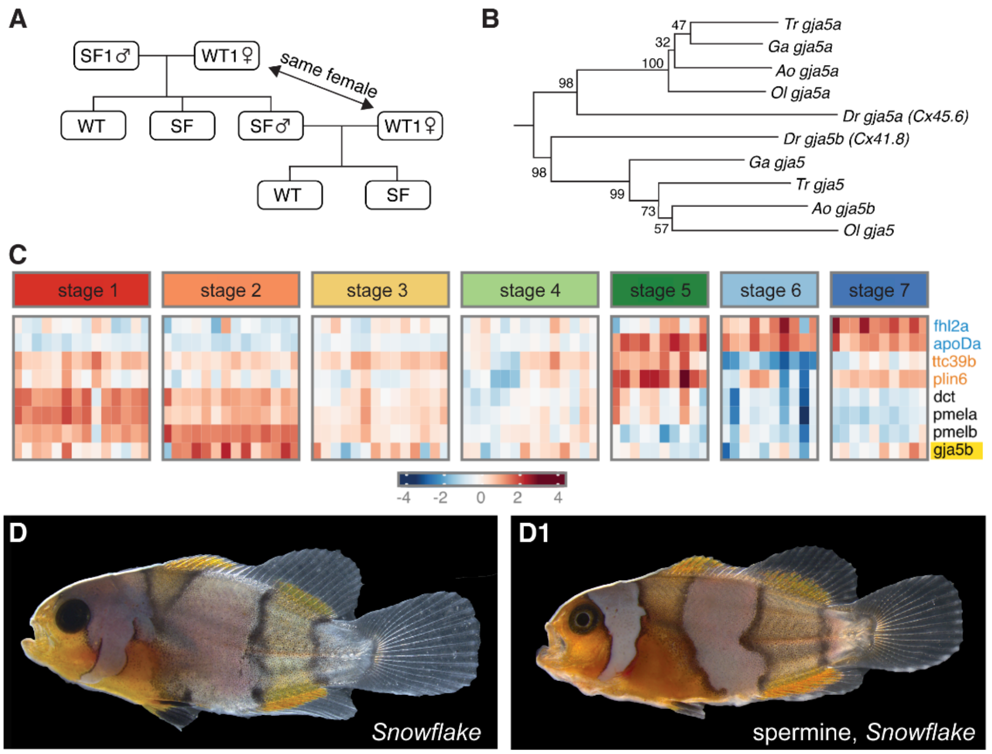

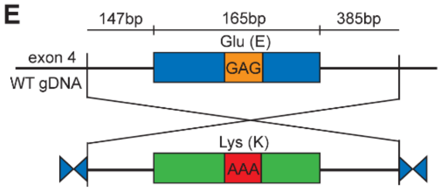

F

| | G | | | CRISPR | A | | | | G | | | CRISPR |
|---|---|---|---|---|---|---|---|---|---|---|---|---|
| | T | G | C | A | T | G | C | A | T | G | C | A |
| WT | 11 (0%) | 89919 (100%) | 35 (0%) | 20 (0%) | 4 (0%) | 11 (0%) | 2 (0%) | 89968 (100%) | 10 (0%) | 89918 (100%) | 19 (0%) | 38 (0%) |
| CRISPR fish WT bar | 5 (0%) | 59478 (99%) | 18 (0%) | 536 (1%) | 3 (0%) | 10 (0%) | 0 (0%) | 60006 (100%) | 5 (0%) | 59475 (99%) | 9 (0%) | 553 (1%) |
| CRISPR fish SF bar | 3 (0%) | 55097 (94%) | 24 (0%) | 3281 (6%) | 3 (0%) | 6 (0%) | 2 (0%) | 55934 (100%) | 2 (0%) | 52749 (94%) | 13 (0%) | 3291 (6%) |

G

| | WT fish | CRISPR fish 1 | CRISPR fish 2 | CRISPR fish 3 | CRISPR fish 4 | CRISPR fish 5 |
|---|---|---|---|---|---|---|
| total reads | 32561 | 34142 | 32920 | 35214 | 35761 | 34790 |
| *gja5b* WT sequence (%) | 32561 (100) | 4097 (12) | 55 (0.1) | 147 (0.4) | 0 (0) | 270 (0.8) |
| *gja5b* total mutant sequences | 0 | 30045 | 32865 | 35067 | 35761 | 34520 |
| *gja5b* short deletion sequences (%) | 0 (0) | 13387 (39) | 16066 (49) | 7883 (22) | 11348 (32) | 18190 (52) |

**Fig. S2. *gja5b* as the gene underlying the *Snowflake* phenotype.**

(**A**) Schematic representation of the backcross for GWAS analysis. (**B**) Phylogenetic tree of the conserved region of *gja5* genes in 5 teleost species indicates that the gene mutated in anemonefish is orthologous to *Cx41.8/gja5b* in zebrafish. Ao – *Amphiprion ocellaris*; Dr – *Danio rerio*; Ga – *Gasterosteus aculeatus*; Ol – *Oryzias latipes*; Tr – *Takifugu rubripes* (**C**) Whole-larvae transcriptomic analysis across the seven larval stages of *A. ocellaris* (*43*) with columns of heat maps representing replicate libraries. Iridophore markers (*fhl2a* and *apoDa*) were expressed at higher levels during later larval stages (stages 5, 6 and 7). Xanthophore markers (*ttc39b* and *plin6*) were expressed throughout larval stages, with an upsurge in stage 5. Melanophore markers (*dct* and *pmela/b*) were highly expressed at the beginning of larval development. The expression of *gja5b* is high initially, peaking at stage 2 and declining thereafter. (**D**) Treatment of pre-metamorphic *Snowflake* fish with spermine (a gap junction inhibitor) does not rescue nor worsen the phenotype. (**E**) Strategy to replace wild-type exon 4 of *gja5b* with exon 4 containing the substitution. (**F**) Results of NextSeq 1000 Illumina sequencing to confirm substitutions achieved by HDR CRSIPR/Cas9 editing. (**G**) Results of NextSeq 1000 Illumina sequencing to confirm *gja5b* mutations induced by NHEJ CRSIPR/Cas9 editing.

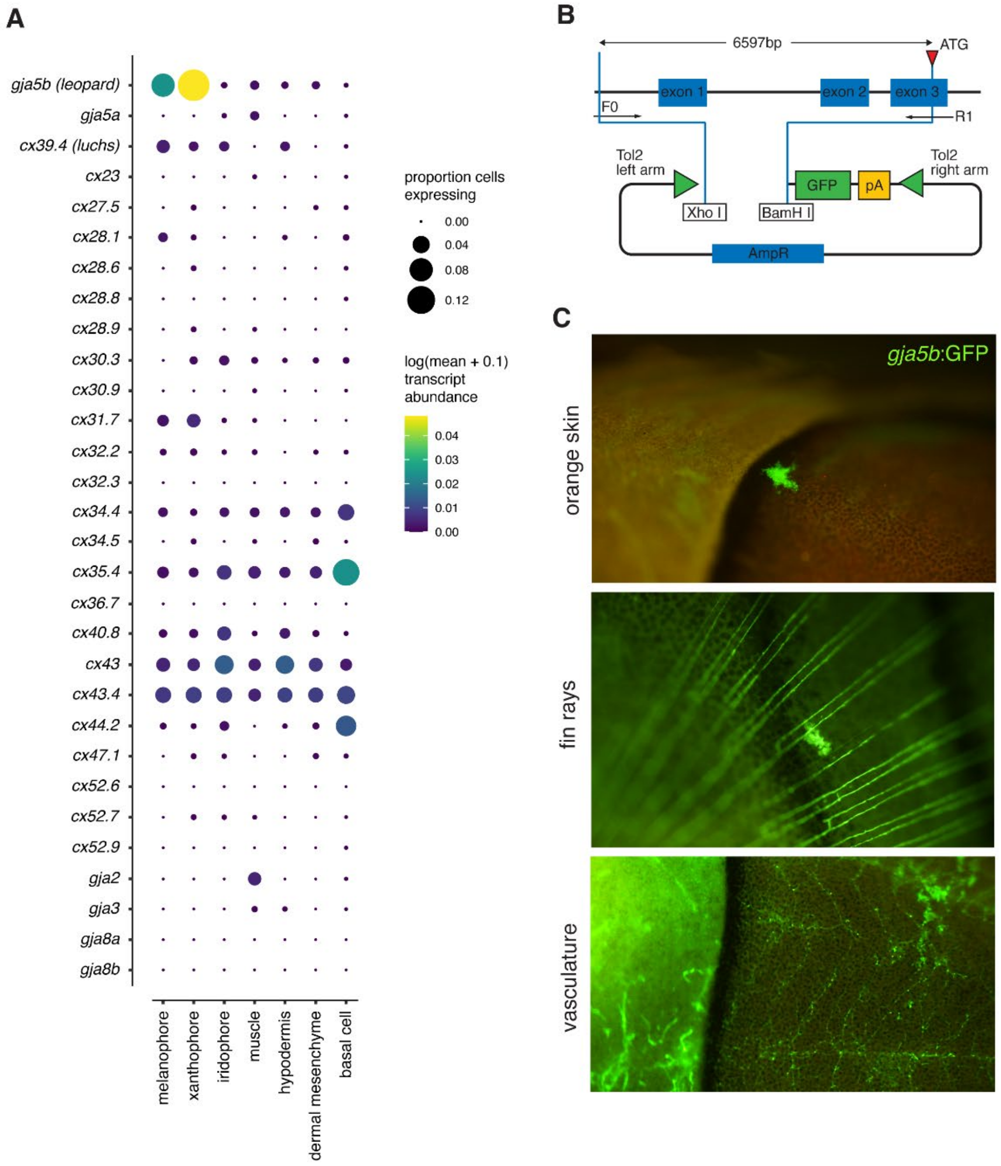

**Fig. S3. *gja5b* expression across species.**
(**A**) In zebrafish, *Danio rerio*, *gja5b* is expressed primarily in melanophores and xanthophores. Dot plot shows relative transcript abundances for gap junction genes detected by single cell RNA-sequencing of zebrafish skin during adult stripe formation, across the three major chromatophore classes and cells of the local tissue environment. Data reanalyzed from (*61*). (**B**) Construct used for transgenesis with transposon, Tol2, to introduce into wild-type fish GFP driven by *gja5b* promoter. (**C**) Besides abundant expression in white bars (Fig. 3B) anemonefish *gja5b*:EGFP was detectable in cells of orange skin, in fin rays, and vasculature.

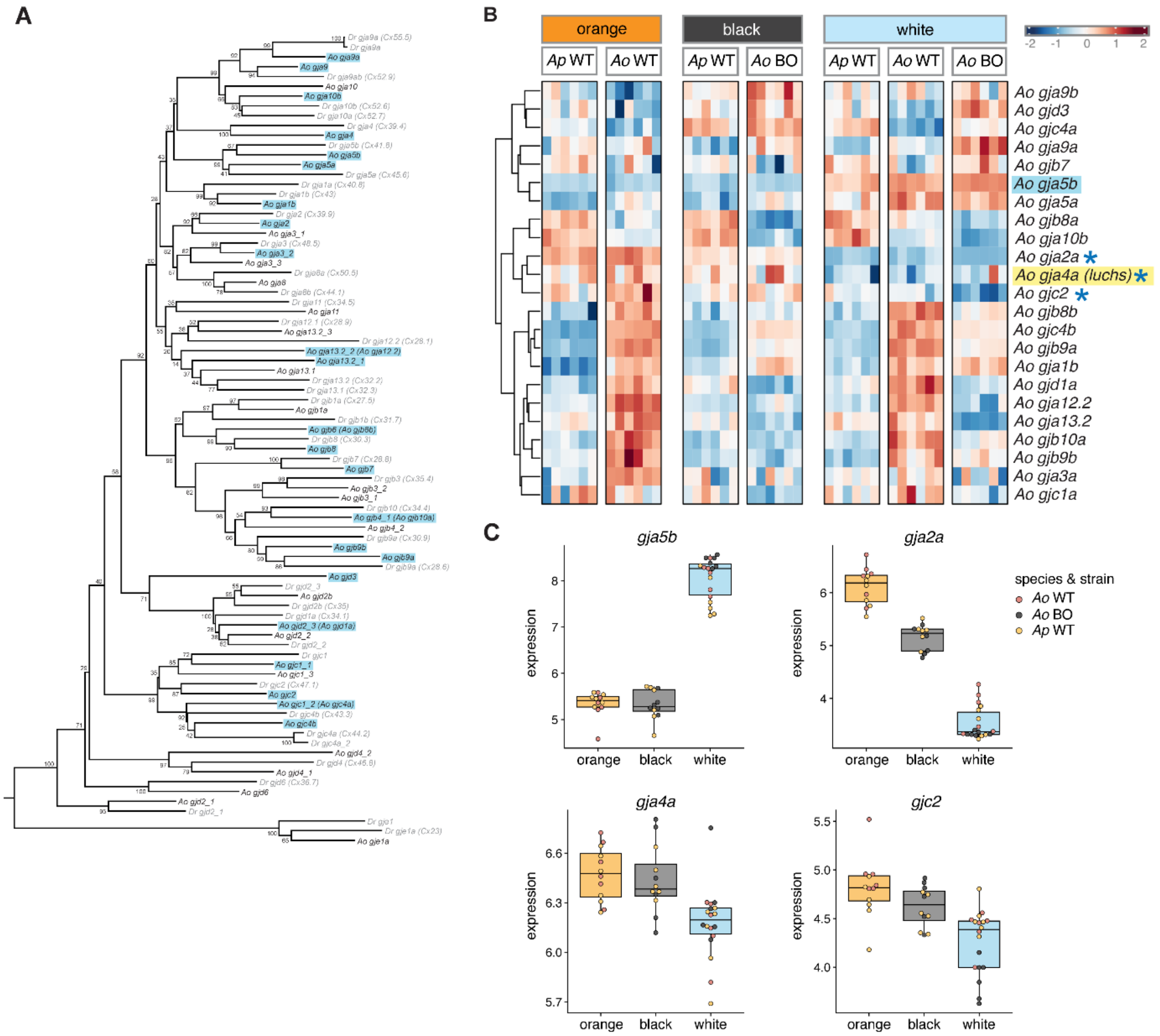

**Fig. S4. Anemonefish gap junction genes and potential for signaling through heteromeric gap junctions.**

(**A**) Reconstruction of evolutionary relationships among gap junction genes of anemonefish (Ao) and zebrafish (Dr), with blue highlighting genes expressed in colored skin regions of anemonefish. (**B**) Heatmaps showing expression of gap junction genes found in color-region specific mRNA sequencing from scales. Scales were collected from *A. ocellaris* wild-type fish (*Ao* WT), *Black A. ocellaris* mutant (*Ao* BO) and *A. percula* wild-type (*Ap* WT). Protein products of the genes marked with an asterisk (*gja2a*, *gja4a*, and *gjc2*) were tested for their ability to form heteromeric channels with Gja5b. *gja2a* transcripts were abundant in xanthophore-enriched orange scales of anemonefish, though the zebrafish orthologue of this gene was expressed primarily in muscle rather than chromatophores (see Fig. S3A). (**C**) Boxplots of gene expression across species, strains and replicate libraries. Abbreviations as above.

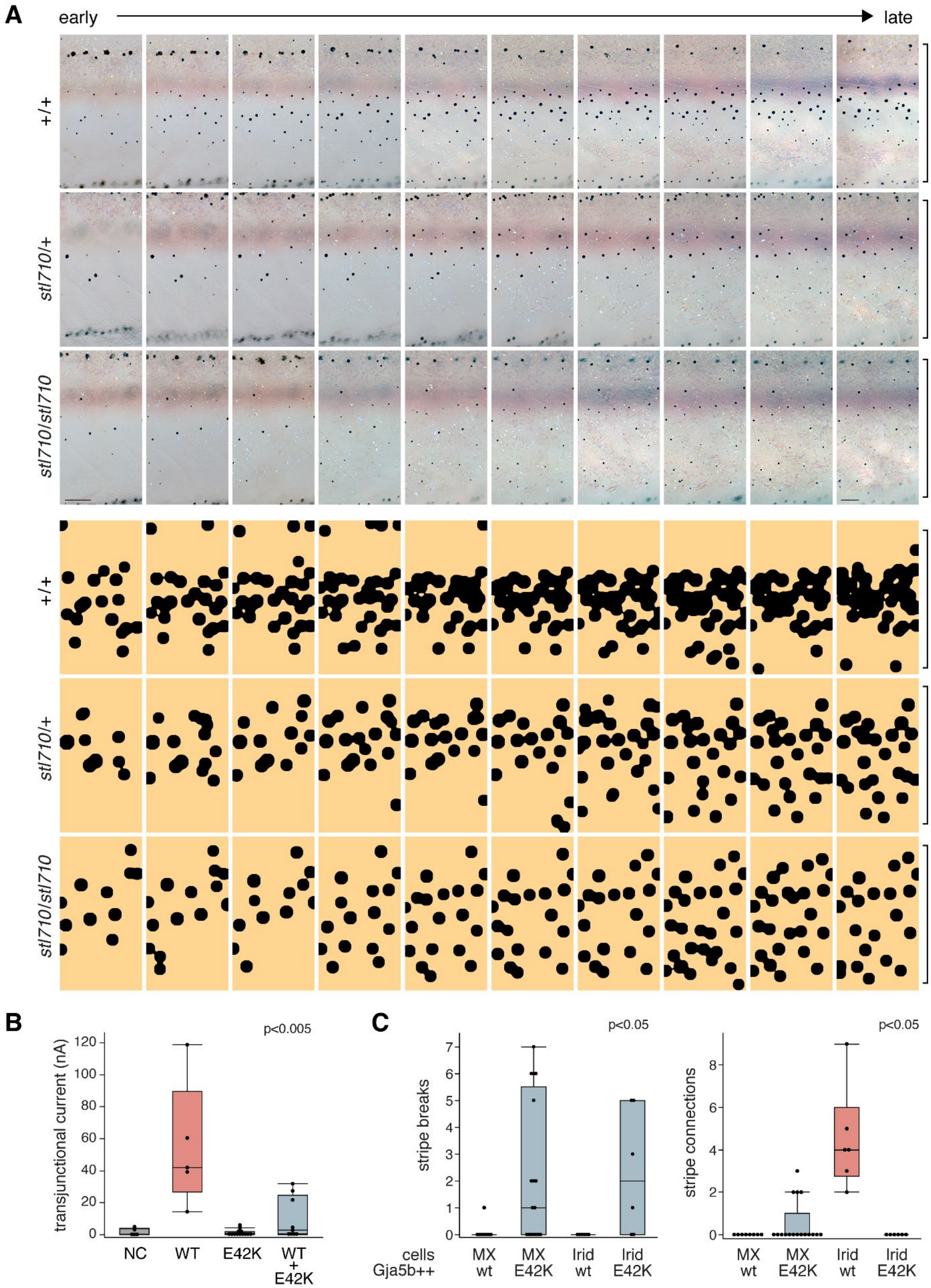
A
early
late
+/+
stl710/+
stl710/stl710
B
p<0.005
transjunctional current (nA)
120
100
80
60
40
20
0
NC
WT
E42K
WT
+
E42K
C
p<0.05
stripe breaks
7
6
5
4
3
2
1
0
cells
Gja5b++
MX
wt
MX
E42K
Irid
wt
Irid
E42K
p<0.05
stripe connections
8
6
4
2
0
MX
wt
MX
E42K
Irid
wt
Irid
E42K

**Fig. S5. An E42K allele of zebrafish *gja5b*.**

(**A**) Ontogeny of pattern in representative fish of indicated genotypes, imaged daily through adult pattern formation. Same ventral regions are shown, with rescaling for growth. Brackets limits at right demarcate positions of horizontal myoseptum (top) and ventral margin of flank (bottom). Upper panel show brightfield imaging of fish treated with epinephrine to contract melanin granules. Lower panels show melanophore positions after pseudocoloring and uniform enlargement of melanin spots to aid visualization of pattern emergence; only newly differentiating adult melanophores are included (*2*). From early to late stages of pattern formation a ventral melanophore stripe becomes increasingly evident in the wild-type (+/+), whereas newly arising melanophores in heterozygotes and particularly homozygotes exhibit a more dispersed arrangement. (**B**) *Xenopus* oocyte assay for zebrafish wild-type Gja5b (pink) and Gja5b with E42K mutation (blue-grey) leading to nearly complete loss of current in homomeric junctions and severely reduced current in heteromeric channels with wild-type. (**C**) Numbers of breaks in stripes, representing expansion of light interstripes, were significantly greater in fish expressing E42K mutant Gja5b in either melanophores and xanthophores (MX) or iridophores (Irid) as compared to fish expressing wild-type Gja5b. Conversely, overexpression of wild-type Gja5b in iridophores led to significantly more connections between dorsal and ventral stripes, a phenotype never observed in wild-type fish. All tests of overall significance by Kruskal-Wallis. Scale bars, 100 mm (A, early at left and late at right).

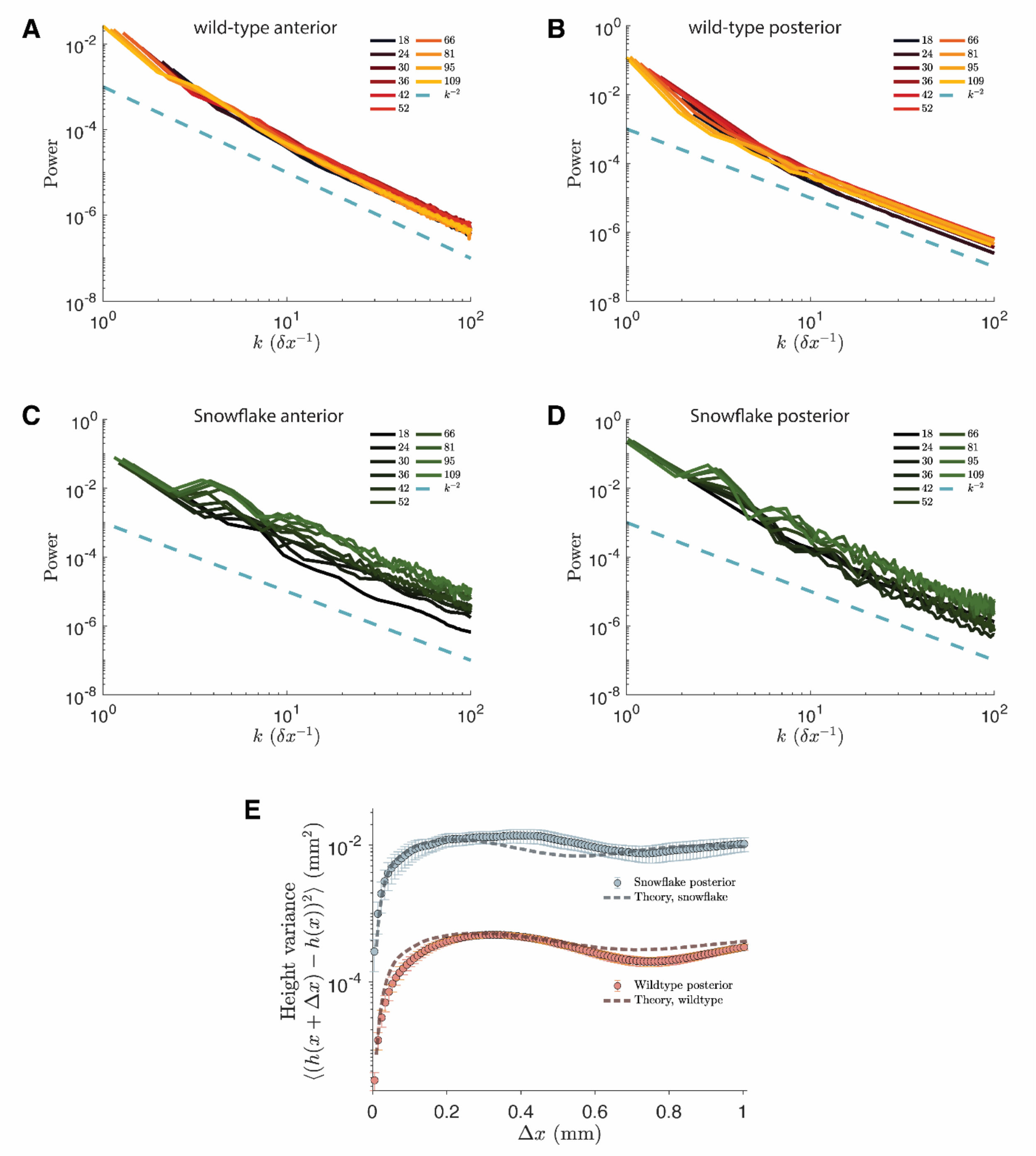

**Fig. S6. Power spectrums of WT and SF bar edges, and posterior bar height-variance.**
(**A**) Average power spectrum of the anterior boundary of the trunk bar in wild-type. Inset legend indicates days-post-hatching. (**B**) Average power spectrum of the posterior boundary of the trunk bar in wild-type. (**C**) Average power spectrum of the anterior boundary of the trunk bar in *Snowflake*. (**D**) Average power spectrum of the posterior boundary of the trunk bar in *Snowflake*. (**E**) Evolution of height variance along the posterior boundary of the trunk bar, comparing experimental data with predictions from the Edwards–Wilkinson model.

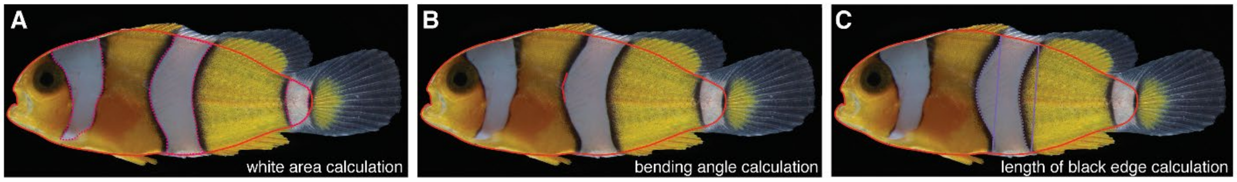

D

| | | | | | | | | | | | | | |
|---|---|---|---|---|---|---|---|---|---|---|---|---|---|
| wild-type gja5b sequence | ACG | GCA | GCC | GAG | TCT | TCG | TGG | GGC | GAC | GAG | CAG | GAA | GAT |
| donor plasmid sequence | ACc | GCt | GCa | AAA | agc | TCc | TGG | GGa | GAC | GAa | CAG | GAg | GAc |

**Fig. S7. Schematic representation of the different metrics calculated to compare *Snowflake* and wild-type siblings and HDR CRISPR donor sequence.**

(**A**) Calculation of white bar areas. (**B**) Calculation of the bending angle. (**C**) Calculation of the length of the black edges. (**D**) Illustration of *gja5b* wild-type coding sequence and oligonucleotide knock-in sequence to confirm successful CRISPR/Cas9 substitution. Red capital letters indicate the desired codon change, while small orange letters indicate the induced silent mutations.

| | | genotype | | |
|---|---|---|---|---|
| gene | phenotype fish | homozygous wildtype | heterozygous | homozygous mutated |
| arfgef3 | wildtype | 19 | 0 | 0 |
| | Snowflake | 1 | 16 | 0 |
| kif20b | wildtype | 18 | 1 | 0 |
| | Snowflake | 2 | 18 | 0 |
| ppp13rca | wildtype | 21 | 1 | 0 |
| | Snowflake | 0 | 18 | 0 |
| LOC111562834 | wildtype | 20 | 0 | 0 |
| | Snowflake | 2 | 13 | 0 |
| LOC111562825 | wildtype | 7 | 2 | 0 |
| | Snowflake | 0 | 11 | 0 |
| gja5b | wildtype | 15 | 0 | 0 |
| | Snowflake | 0 | 30 | 0 |
| znf318 | wildtype | 16 | 1 | 0 |
| | Snowflake | 0 | 19 | 0 |

**Table S1: Potential candidate genes other than *gja5b* could be excluded using Sanger sequencing.**

| variant type | number of SNVs | frequency of SNVs |
|---|---|---|
| intergenic region | 981 | 53.67% |
| intron variant | 580 | 31.73% |
| downstream gene variant | 79 | 4.32% |
| upstream gene variant | 129 | 7.06% |
| 3' UTR variant | 22 | 1.20% |
| 5' UTR variant | 13 | 0.71% |
| synonymous variant | 14 | 0.76% |
| missense variant | 10 | 0.55% |
| total | 1828 | 100% |

**Table S2: Detailed single nucleotide variant analysis of chromosome 16.**

| | primer name | primer sequence (5'-3') |
|---|---|---|
| Snowflake confirmation | gja5-gt-F2 | CAGATAAGACCCGACAGGAATGTAATCC |
| | gja5-gt-R2 | CTGTAAAATAGGATTATAGGTATAGGTGAAAAGTG |
| | arfgef3-gt-F1 | GAAGCAGCTGTGTCCGTCTCTGG |
| | arfgef3-gt-R1 | CTCCTTCATGATGCGGATGGCC |
| | kif20b-gt-ex29-F3 | CTGTTCAGCTTTCTGTTGGGTGC |
| | kif20b-gt-ex29-R3 | CTCCCGCTGTGTAATTTAACCTCAC |
| | Kif20b-g-exon20-F1 | GAGGCTTTGACTGCCCTCGAAAAGG |
| | Kif20b-g-exon20-R1 | CTTGTGCAGCTTTCTCTGTCTCGATGGTC |
| | Kif20b-g-exon 22-F2 | GTGATCTTCCCATGAGCAGAAGACAACCG |
| | Kif20b-g-exon 22-R2 | GAGGGTCCAATTCAACCCGTGGG |
| | LOC111562825-gt-start | ATGGAGGGACGTTCTGCTGTTTGGC |
| | LOC111562825-gt-R2 | CACTCTGAATCCTGTTGTTGATCGCC |
| | LOC111562834-gt-F2 | GCTCTGATCTCTGCTGAAGCCCAG |
| | LOC111562834-gt-R2 | CAGATTGTCCACAGGTTCACTGTCTTTCTC |
| | ppp1r3ca-gt-F1 | GATCTGCACAGGGCCAAGAACCG |
| | ppp1r3ca-gt-R1 | GCTGTCGTTATTGTCCCAGAAGGTCTG |
| | znf318-gt-F1 | CCAAAGATGACAGCGACAGTAAGGCC |
| | znf318-gt-R1 | CCCAGAAACTCAGACTACATGTGGGATAC |
| Xenopus cRNA | gja5b-F | aaatttgaattcgccaccATGGCCGACTGGAGTCTGCT-3 |
| | gja5b-R | ttttgcggccgcCTAAACTGAAAGGTCGTCTGCTCTG |
| | gja2-F | aaaagaattcgccaccATGGCAGACTGGAACCTGTTGGGG |
| | gja2-R | ttttgcggccgcTCACACCTTCAGGTCGTCCAGCCTG |
| | gja4-F | aaaagaattcgccaccATGTCCAGAGCTGACTGGTCCTTC |
| | gja4-R | ttttgcggccgcTCATACATAATGTTTTTGTTTTG |
| | gjc2-F | aaaagaattcgccaccATGAGCTGGAGCTTTCTCACTCGC |
| | gjc2-R | ttttgcggccgcCTAGATCCAGACAGAGGTCTTTCC |
| | cx41.8-F | TAGGCACGGCTGCTAAATCCTCGTGGGGGGACGAGCAGG |

| | | |
|---|---|---|
| | cx41.8-R | CCACGAGGATTTAGCAGCCGTGCCTAGTACCAAGATCCG |
| CRISPR related | Ao-gja5-HAL-XhoI-F | GAGGctcgagCAGATAAGACCCGACAGGAATGTAATCC |
| | Ao-gja5-SF-R | TCGTCTCCCCAGGAGCTTTTTGCAGCGGTACCGAGAACCAGGATCCG |
| | Ao-gja5-mut-F | AGCTCCTGGGGAGACGAACAGGAGGACTTTAACTGTGACACCGAACAGC |
| | Ao-gja5-HAR-SpeI-R | GAGGactagtCTGTAAAATAGGATTATAGGTATAGGTGAAAAGTG |
| | Ao-gja5bPro-F0 | ATCACTTGGGCCCGGCTCGAGTGACTGATGGAACTCTTCATGGTC |
| | Ao-gja5bPro-R1 | ATGGTGGCGACCGGTGGATCAACAGTTATAGATCAGGTCAGAGG |
| | Ao-gja5-KI-check-F2 | CTTGTAAGTATTAAGGTGAAGATTCTACAT |
| | Ao-KI-check-R | GTCCTCCTGTTCGTCTCCCCAGGAGCT |
| | TruSeq-F | ACACTCTTTCCCTACACGACGCTCTTCCGATCT |
| | | GGAGGTCTTGATATTTCTTGATTTATTTCCTC |
| | TruSeq-R | GTGACTGGAGTTCAGACGTGTGCTCTTCCGATCT |
| | | CGGTCGTAACAAACGTTCTCACAGCCTGGC |

**Table S3: Primer details.** Small letters indicate added restriction enzyme sites.

**Movie S1:** Sequential days in development of a wild-type zebrafish showing emergence of adult stripes. Images were rescaled and aligned to control for growth over 30 d.

**Movie S2:** Sequential days in development of a *gja5b*$^{stl710}$ homozygous mutant zebrafish (sibling to fish of Movie S1) showing ontogeny of relatively uniformly dispersed melanophores. Images rescaled and aligned to control for growth over 32 d.
